# An early cell shape transition drives evolutionary expansion of the human forebrain

**DOI:** 10.1101/2020.07.04.188078

**Authors:** Silvia Benito-Kwiecinski, Stefano L. Giandomenico, Magdalena Sutcliffe, Erlend S. Riis, Paula Freire-Pritchett, Iva Kelava, Stephanie Wunderlich, Ulrich Martin, Greg Wray, Madeline A. Lancaster

**Affiliations:** MRC Laboratory of Molecular Biology, Cambridge Biomedical Campus, Francis Crick Avenue, Cambridge, CB2 0QH; Department of Applied Mathematics and Theoretical Physics, University of Cambridge, Wilberforce Road, Cambridge, CB3 0WA; Leibniz Research Laboratories for Biotechnology and Artificial Organs (LEBAO), REBIRTH-Research Center for Translational and Regenerative Medicine, Hannover Medical School, 30625 Hannover, Germany; Biomedical Research in Endstage and Obstructive Lung Disease (BREATH), Member of the German Center for Lung Research (DZL), 30625 Hannover, Germany; Department of Biology, Duke University, Biological Sciences Building, 124 Science Drive, Durham, NC, 27708, USA

**Keywords:** Brain evolution, organoids, cell shape, morphogenesis, neuroepithelium

## Abstract

The human brain has undergone rapid expansion since humans diverged from other great apes, but the mechanism of this human-specific enlargement is still unknown. Here, we use cerebral organoids derived from human, gorilla and chimpanzee cells to study developmental mechanisms driving evolutionary brain expansion. We find that the differentiation of neuroepithelial cells to neurogenic radial glia is a protracted process in apes, involving a previously unrecognized transition state characterized by a change in cell shape. Furthermore, we show that human organoids are larger due to a delay in this transition. Temporally resolved RNA-seq from human and gorilla organoids reveals differences in gene expression patterns associated with cell morphogenesis, and in particular highlights *ZEB2*, a known regulator of epithelial-mesenchymal transition and cell shape. We show, through loss- and gain-of-function experiments, that *ZEB2* promotes the progression of neuroepithelial differentiation, and its ectopic overexpression in human is sufficient to trigger a premature transition. Thus, by mimicking the nonhuman ape expression in human organoids, we are able to force the acquisition of nonhuman ape architecture, establishing for the first time, an instructive role of neuroepithelial cell shape in human brain expansion.

## Introduction

One of the most distinctive features of humans as a species is our enlarged brain. While brain expansion is a hallmark of primate evolution in general, larger brains have been particularly selected for along the human lineage. The human brain is the largest of all primates and is roughly 3-fold larger than that of our closest living relatives, the chimpanzee and gorilla (Herculano-Houzel, 2012). Most of our understanding of the principles governing mammalian brain development comes from studying the mouse, and although the stages and cytoarchitectural features of brain development are broadly conserved amongst mammals, the end-product is substantially different in humans compared to the mouse, most notably in size, with more than a 1,000-fold increase in total neuron number (Herculano-Houzel et al., 2006). Thus, a number of studies have examined differences in brain development between human, mouse and other distantly related model organisms (Bae et al., 2015; Lui et al., 2011; Sousa et al., 2017), highlighting divergence in neural progenitor behaviour (Florio et al., 2015), neurogenesis (O’Neill et al., 2018), and cytoarchitecture (Dehay et al., 2015; Fietz et al., 2010; Hansen et al., 2010). While much has been learned about the genetic and cell biological mechanisms governing the evolutionary divergence from rodents, human-specific changes compared to other apes are less well understood.

Comparative studies have led to a number of observations that might help explain the human-specific difference in brain size. One such observation is that the human brain appears to be a scaled-up primate brain, meaning that it conforms to primate scaling rules, and thus contains the expected number of neurons for a primate brain of its size (Herculano-Houzel, 2009). Similarly, comparative neuroanatomical studies of adult ape brains have revealed a general increase in size, without a disproportionate increase in particular layers of the cerebral cortex (De Sousa et al., 2010). Given that cortical layers are generated successively over time, these findings would suggest that human-specific differences governing brain size take place prior to their generation, even before the onset of neurogenesis.

Neurogenesis begins when precursor cells called neuroepithelial (NE) cells make the key transition to neurogenic radial glia (RG). Before this transition, NE cells exhibit a columnar morphology and divide in a symmetric proliferative manner. This proliferation results in an exponential expansion, and in the neocortex this expansion leads to a ballooning out of the tissue and enlargement of the ventricles. In humans, this ballooning is particularly pronounced (Bayer et al., 1993). NE cells are strongly epithelial in character and held together near their apical surface by tight junctions and adherens junctions. In rodents and other model vertebrates, it is well-documented that the transition to neurogenic RG cells involves a loss of epithelial features, such as tight junctions, a thinning and elongation of the bipolar processes extending apico-basally from the cell body, and a switch to asymmetric cell division with one daughter remaining a RG, and the other becoming more differentiated and leaving the ventricular zone (Götz and Huttner, 2005). This switch in cell fate thus leads to a change in expansion from tangential to radial, with additional cells being added to the more basal layers, rather than along the apical surface of the ventricles.

This model of NE-to-RG conversion is largely based on studies in mouse, where between E9 and E10 cells rapidly undergo this switch characterised by simultaneous changes in cell morphology, cell-adhesion, and molecular identity (Kriegstein and Alvarez-Buylla, 2009). However, it remains entirely unknown how this process occurs in apes. It has long been hypothesised that changes in NE behaviour could lead to an expansion of the neocortical primordium in the tangential dimension and the establishment of a larger protomap, which might have been important in human brain expansion (Rakic, 1988, 1995). Several lines of evidence support this; firstly, the human forebrain is already larger than mouse and macaque at the onset of neurogenesis, indicating differences in NE expansion (Rakic, 2007). Secondly, while human and chimpanzee brains are comparable in mean cortical thickness, there is a more than 3-fold difference in cortical surface area (Donahue et al., 2018), further pointing towards early changes in tangential expansion as being key to later size differences between apes. However, without being able to examine and manipulate early NE behaviour in apes and humans, these observations have remained correlative.

The ability to generate brain organoids from induced pluripotent stem cells (iPSCs) has enabled researchers to study a variety of neurodevelopmental processes that were previously inaccessible (Lancaster et al., 2013; Kadoshima et al., 2013; Pasca et al., 2015; Quadrato et al., 2017). Recently, a number of single-cell transcriptomic studies have started tackling evolutionary differences across primates, highlighting human-specific patterns of gene expression (Kanton et al., 2019; Mora-Bermúdez et al., 2016; Pollen et al., 2019). Human organoids appear to develop and mature at a slower pace relative to chimpanzee and macaque, with neurons in the latter two species showing an increased expression of genes related to neuronal maturation (Kanton et al., 2019). Comparative analysis of human, chimpanzee and macaque showed that neurogenesis commences earlier and proceeds faster in macaque (Otani et al., 2016). A further comparison between human, chimpanzee and macaque organoids, revealed that human RG cells display a longer mitotic phase (Mora-Bermúdez et al., 2016), which has previously been linked to RG cells that can undergo proliferative divisions, rather than asymmetric neurogenic divisions. This suggests that even after the onset of neurogenesis, human RGs may favour proliferative divisions over neurogenic divisions for a longer period, which is consistent with a larger brain. Similarly, NOTCH2NL, a family of human-specific paralogs of NOTCH2, was shown to promote RG expansion over neurogenesis, with *NOTCH2NL* knockout organoids displaying premature neurogenesis and neuronal maturation (Fiddes et al., 2018, Suzuki et al., 2018). Finally, an alternative approach was the use of transgenic mice carrying either the human or the chimpanzee region of a divergent genetic locus (Boyd et al., 2015). Here the authors demonstrated an effect of the human sequence on expression of Fzd8, resulting in an accelerated cell cycle of RG cells and increased neocortical size. Although these studies point to important differences in terms of RG fate decisions and maturation rates, which likely have implications for brain size determination, all of these comparisons have been performed at stages where neurogenesis is already underway. Thus, any differences would affect neuron types generated afterwards, and would therefore be expected to lead to a disproportionate increase in only those later-born cell types.

Given the neuroanatomical comparisons that point to an early expansion and to a universal increase in neurons, it would be highly informative to examine early human and ape NE at a stage when neurons have not yet been generated. Furthermore, extending comparative analysis to include other great apes beyond chimpanzee would help to identify true human-specific cellular adaptations (rather than chimpanzee-specific differences) involved in brain size determination. Finally, there is now a wealth of data generated by comparative genetic analyses (Dorus et al., 2004; Enard, 2016; Gordon et al., 2016; Pääbo, 2014; Pollard et al., 2006; Prüfer et al., 2012), but functional studies will be essential in understanding how these genetic changes translate into cellular phenotypes that explain evolutionary differences.

In an effort to dissect the cellular and tissue level changes occurring during human and non-human ape brain development, we examined human, chimpanzee and gorilla organoids, focusing on NE development prior to the onset of neurogenesis. We found that human organoids displayed differences in tissue architecture at early stages of NE expansion, before the onset of neurogenesis, with a notable enlargement of the ventricular apical surface. Through careful cellular analysis, we found that unlike the rapid differentiation of NE to RG cells described in mouse, the transition in human and ape is a gradual process, taking place over the course of several days and involving an intermediate cell morphology state that we have named transitioning NE (tNE) cells. Importantly, we found that ape NE cells make this transition more rapidly than human, but in both the cell shape change occurs before the change in cell identity and the onset of neurogenesis. To identify the molecular mechanism underlying these differences, we performed highly time-resolved deep sequencing analysis and identified differential expression dynamics of the zinc-finger transcription factor *ZEB2*. Through loss- and gain-of-function experiments with *ZEB2* we show that this gene is a driver of the NE to RG transition. Importantly, we demonstrate that elevating *ZEB2* levels in human organoids, similar to gorilla, is sufficient to trigger precocious loss of epithelial characteristics and gain of tNE cell morphologies, which leads to smaller tissue size and mimics the ape phenotype. Thus, our data suggest that in humans a delayed onset of *ZEB2* expression extends the NE stage, compared to ape, and may be a major contributor to neocortical expansion in humans.

## Results

### Human telencephalic organoids show differences in tissue architecture prior to neurogenesis

To investigate evolutionary differences in brain development and brain size determination between apes, we generated cerebral organoids from human, gorilla and chimpanzee cell lines. Importantly, we optimised the protocol so that an identical approach of neural induction, expansion and growth of organoids with a telencephalic identity could be used for all species (Figure S1A) with highly comparable resulting tissues (Figure 1A). This was vital to be able to attribute any differences to species-specific divergence and not protocol variations. We did, however, notice that prior to neural induction, chimpanzee embryoid bodies (EBs) seemed to progress faster and had to be transferred to neural induction 2 days earlier than human and gorilla EBs, potentially reflecting the shorter gestational period of chimpanzees relative to the two other apes (Ardito, 1976).

**Figure 1.**
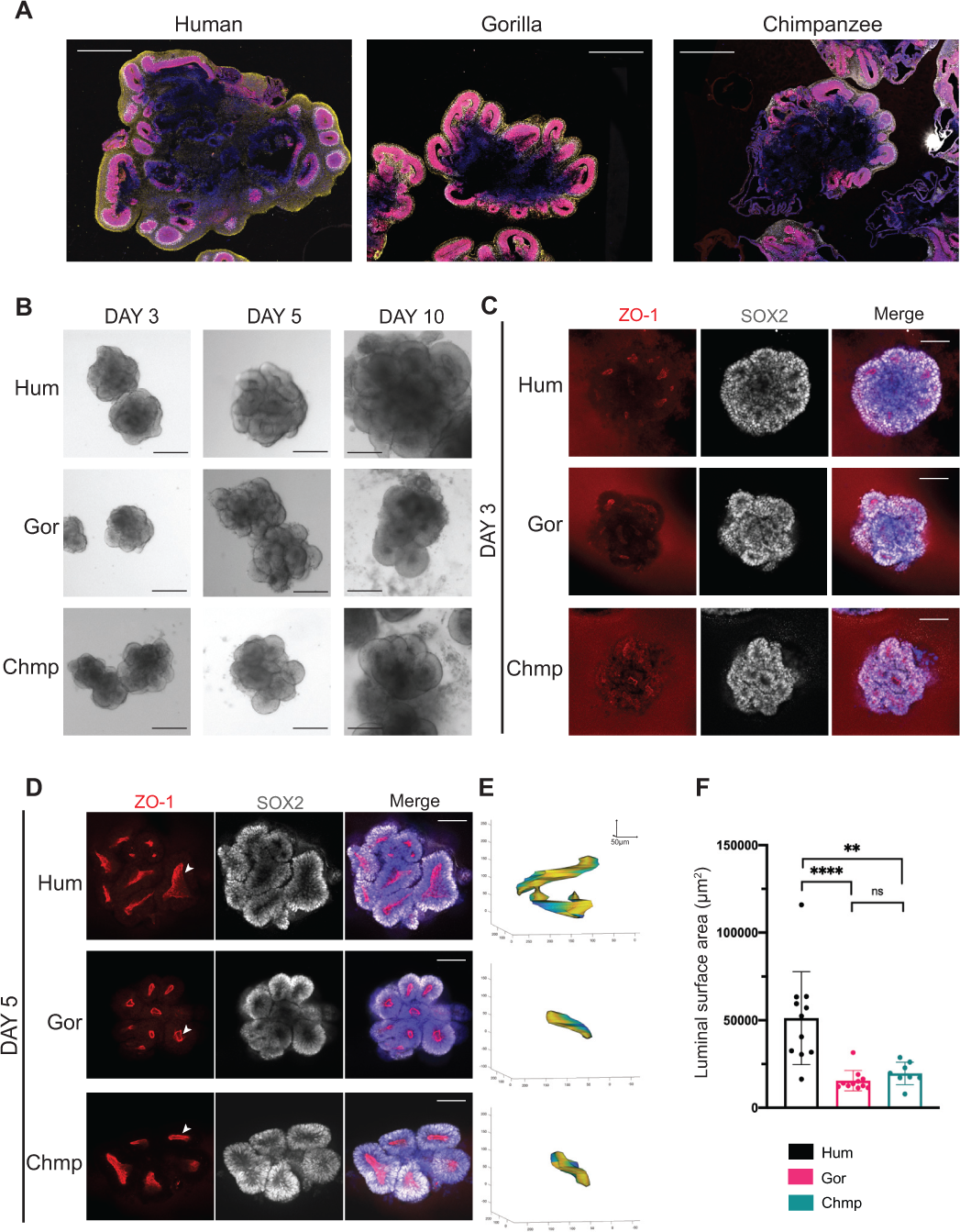
Human telencephalic organoids are larger and exhibit larger apical lumens than organoids from other apes. **A.** 5-week organoids stained for neural progenitor marker SOX2 (red), dorsal telencephalic/intermediate progenitor marker TBR2 (grey),neuronal markers TUJ1 (human) and HuCD (gorilla) in yellow, and DAPI (blue) showing human (H9) derived organoids become larger in overall size than gorilla (G1) and chimpanzee (Chmp) organoids. Scale bar: 1mm. **B.** Brightfield images of ape organoids at day 3, 5 and 10 showing human (H9, top panel) differences in tissue architecture arising at day 5 with more elongated neuroepithelial buds versus more circular buds seen in nonhuman ape organoids (G1, Chmp, middle and bottom panel). Scale bar: 200 μm. **C, D.** Representative immunofluorescence images of the center of whole mount human (H9), gorilla (G1), and chimpanzee (Chmp) organoids with staining for ZO1 and SOX2 showing polarized neural progenitor cells organized around ZO1 positive apical lumens at day 3 (**C**) and day 5 (**D**). Note the appearance of less rounded ZO1 positive apical lumens (arrowheads) in human organoids relative to nonhuman ape organoids. DAPI is in blue. Scale bar: 100 μm. **E.** 3D Matlab reconstructions of apical lumens of day 5 organoids showing an example of the more convoluted shapes found in human versus chimpanzee and gorilla derived organoids. Figures are a representative example of 3D reconstructions performed on the largest apical lumen per organoid. Values on the axes are in μm. Luminal surface area of reconstructed examples: human (H9) = 63,394 μm^2^; gorilla (G1) = 15,146 μm^2^; chimpanzee (Chmp) = 19,730 μm^2^. **F.** Quantification of the surface area of the largest apical lumen per day 5 organoid reveals significantly expanded luminal surface areas in human versus nonhuman apes. Mean luminal surface area: human (H9) = 51,243 μm^2^; gorilla (G1) = 15,437 μm^2^; chimpanzee (Chmp) = 19,632 μm^2^. *P<0.05, ** P = 0.0012, **** P<0.0001, one-way Anova and post-hoc Tukey’s multiple comparisons test, n (H9 and G1) = 11 organoids from 5 independent batches, n (Chmp) = 8 organoids from 2 independent batches, error bars are S.D.

We immediately noticed that human organoids were consistently larger than gorilla and chimpanzee organoids (Figure 1A). The presence of TBR2+ intermediate progenitors (Figure S1B) confirmed primarily dorsal telencephalic identity in organoids across all three species. Despite the larger overall size, individual cortical lobules exhibited similar thickness and relative proportion of progenitor and neuron types (Figure S1B) suggesting the overall size differences might be due to earlier differences in the establishment and expansion of founder progenitor cell populations.

In order to decipher the exact stage at which size differences become apparent, we examined the development of organoids from the earliest stages of neural tissue formation. After Matrigel embedding, organoids of all species displayed well-formed NE buds (Figure S1C) of similar sizes. Staining for ZO1, a tight junction protein that marks the apical surface of polarized epithelia (Ando-Akatsuka et al., 1999; Medelnik et al., 2018) revealed similar tissue architecture across species, with SOX2+ neural progenitor cells organized in a narrow layer around ZO1+ apical lumens (Figure 1C). At day 5, however, we observed clear differences in tissue architecture between human organoids and those of the other apes. Whilst gorilla and chimpanzee organoids presented more rounded circular neural buds, human organoids showed more elongated neural buds (Figure 1B, S1C). ZO1 staining revealed that the more circular buds of gorilla and chimpanzee organoids corresponded to more rounded apical lumens, whereas human organoids consisted of lumens with more convoluted shapes (Figure 1D, S1D). Using a custom image analysis pipeline, we traced out and reconstructed ZO1+ apical lumens in 3D, revealing species-specific differences in luminal shape and size (Figure 1E). Quantification revealed consistently and significantly enlarged luminal surface area in human organoids compared with other apes (Figure 1F, S1E).

We next sought to focus on the cell biological mechanisms that might explain these differences. In order to ensure a robust comparative analysis, we decided to focus on human and gorilla organoids, made from two cell lines each, as they exhibited identical developmental progression (Figure S1A) during all stages prior to the onset of size differences at day 5. Given current knowledge of early cortical development, the most likely explanation for the expanded lumens observed in human organoids would be a delay, relative to gorilla, in the switch from symmetrically expanding NE cells to neurogenic RG cells. We therefore tested whether at this stage, gorilla organoids had already switched to generating neurons, by staining for neurogenic TBR2+ intermediate progenitors and DCX+ newborn neurons. Surprisingly, in both human and gorilla organoids, neither cell type was present at day 5, only becoming clearly evident between days 10 and 15 (Figure 2A, Figure S2A). This suggests that a faster neurogenic switch in gorilla compared to human does not explain the difference in size of cortical tissues.

**Figure 2.**
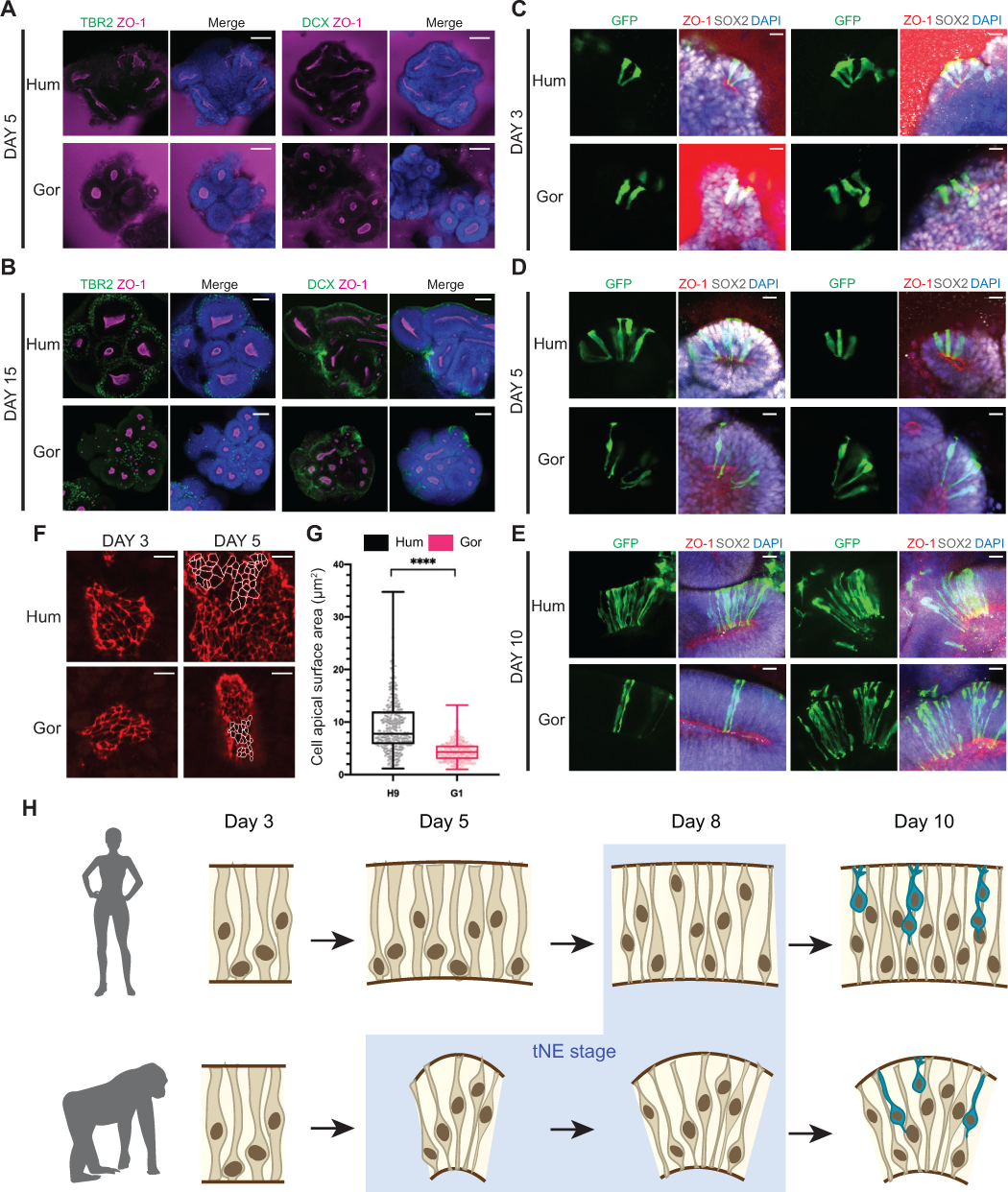
Human NE cells exhibit delayed cell shape transition. **A,B.** Representative immunofluorescence images through whole mount human (H9) and gorilla (G1) derived organoids for early indicators of neurogenesis, DCX (newborn neurons) and TBR2 (intermediate progenitors), show that neurogenesis has not started at day 5 (**A**) but is underway by day 15 (**B**). DAPI is shown in blue. Scale bar: 100 μm. **C-E.** Representative immunofluorescence images of the morphology of neural progenitor cells (SOX2+), polarized around apical (ZO1+) lumens, revealed by sparse labelling with viral GFP. **C.** Day 3 cells are columnar and exhibit the classical NE shape. **D.** Day 5 human (H9) cells still appear columnar, whereas gorilla (G1) cells show a thinning of apicobasal processes consistent with tNE morphology. **E.** Day 10 cells appear RG-like in both species with elongated and narrowed apicobasal processes. Scale bar: 20 μm. **F.** Immunofluorescent staining for ZO1 on the surface of apical lumens showing the apical surface areas of individual progenitor cells at day 3 and 5 in organoids derived from both species (H9, G1). Perimeters of some individual progenitor cells of day 5 organoids are delineated in white highlighting the more constricted apical surface seen in gorilla organoids. Scale bar: 10 μm. **G.** Quantification of the surface area of individual neural progenitor cells of day 5 organoids show significantly smaller apical surface sizes of gorilla (G1) cells compared to human (H9). Measurements were performed on delineated ZO1 cell perimeters as demonstrated in Figure 2F. Mean apical surface area/cell: human (H9) = 9.39μm^2^, gorilla (G1) = 4.48μm^2^. Mann-Whitney U = 18471, **** P<0.0001, two-tailed, n (H9) = 341 cells from 8 organoids from 2 independent batches, n (G1) = 321 cells from 9 organoids from 2 independent batches, error bars are min-max values, dots on the boxplot represent individual cells. **H.** Schematic summarizing the morphological changes in neural progenitor cells observed in human and gorilla organoids. Progenitor cells of both species undergo a gradual transition from NE to tNE to RG-like shapes. Human cells maintain columnar NE-characteristics for a longer period while gorilla cells show tNE morphologies (blue background) earlier than human.

### The NE to RG switch involves a transitioning NE cell morphotype that is delayed in human

Another possible explanation for the difference in tissue morphology may be a difference in cell shape. We therefore examined cell morphology of SOX2+ neural stem cells using sparse viral labelling with GFP. At day 3, labelled cells of both species exhibited a wide, columnar shape characteristic of NE cells (Figure 2C, S2B). At day 5, however, when the differences in tissue architecture first become apparent, we observed a clear difference in the shape of labelled cells between human and gorilla (Figure 2D, Figure S2B). Whilst in human organoids the majority of cells still appeared wide and columnar in shape, in gorilla, cells exhibited more constricted apical processes than their human counterparts. By day 10, labelled cells in both species exhibited the typical elongated, narrow shape of RG cells (Figure 2E). These findings suggest that the transition from NE to RG involves a change in cell shape before the switch to neurogenic fate. Furthermore, this elongated transition state, herein referred to as transitioning NE (tNE) cells, appears delayed in human organoids.

To quantify these differences in cell shape, we measured the apical surface area of individual progenitor cells using ZO1 staining, which delineates apical edges at the lumen (Figure 2F). The apical surface of cells from day 3 organoids were large and indistinguishable between human and gorilla (Figure 2F). Between day 3 and 10, when cells have fully transitioned from NE-like to RG-like morphologies in both species, the apical surface of progenitors reflected this transition, decreasing 7-fold in size (Figure S2B-E). However, the transition was accelerated in gorilla, where already by day 5 the apical surface of cells was significantly more constricted compared to human (Figure 2F), with a mean surface area roughly half that of human cells (Figure 2G, S2F). These results overall reveal a more rapid switch to apically constricted tNE morphologies in gorilla compared with human (Figure 2H).

We also tracked individual NE cells by live imaging in order to see how cell shape differences between species might arise before day 5 (Movie S1, S2). As previously observed in human NE cells (Subramanian et al., 2017), we observed a transient loss of the basal process during NE cell division in both human and gorilla. Following the division, NE daughter cells in both species initially displayed a thinner apical process than the parent cell. However, the apical processes of the human NE daughter cells gradually began to widen again, whereas gorilla NE cells did not regain apical thickness after division. This suggests an active reacquisition of a wide, columnar morphology in the human at this stage, whereas gorilla had already transitioned to the elongated tNE morphotype.

### RNA-seq analysis captures dynamics of gene expression across multiple early time points

In order to identify what factors might be controlling these changes in cell shape, we examined the transcriptome profile of human and gorilla organoids. While there are a number of published cerebral organoid single cell RNA-seq datasets, including from human and chimpanzee (Kanton et al., 2019; Mora-Bermúdez et al., 2016; Pollen et al., 2019), those collected to date focus on only a few time points, which fail to capture the specific NE to RG transition. Therefore, since we were interested in identifying high confidence differentially expressed genes, which requires high coverage sequencing, and because these early stages are less heterogeneous in terms of cellular make-up, we decided to perform bulk RNA-seq. Furthermore, because we wanted to capture the temporal dynamics during the transition, this approach enabled us to assay a large number of organoids across many time points and replicates. In total, we performed RNA-seq of 42 samples of approximately 9,000 organoids, collected from day 0, corresponding to pre-neurulation tissue, up until day 25, corresponding to fully committed neurogenesis (Figure 3A). In order to compare expression of genes shared between both species, TPMs were calculated using a filtered list of annotated genes for which an orthologous gene was present in both species (Figure S3A). Principal component analysis of the samples showed biological replicates grouping together, highlighting the robustness and reproducibility of the protocol (Figure S3B). At the same time, it also showed samples separated by species along the first principal component, followed by grouping by time along the second and third principal components. PC1 was likely driven by technical artifacts arising from read mapping and annotation differences coming from the reference genomes of the two species. Therefore, in order to normalize within each species and at the same time focus more on differences in gene expression dynamics rather than absolute levels, the expression of each gene was normalized over time by z-scaling across the entire time course.

**Figure 3.**
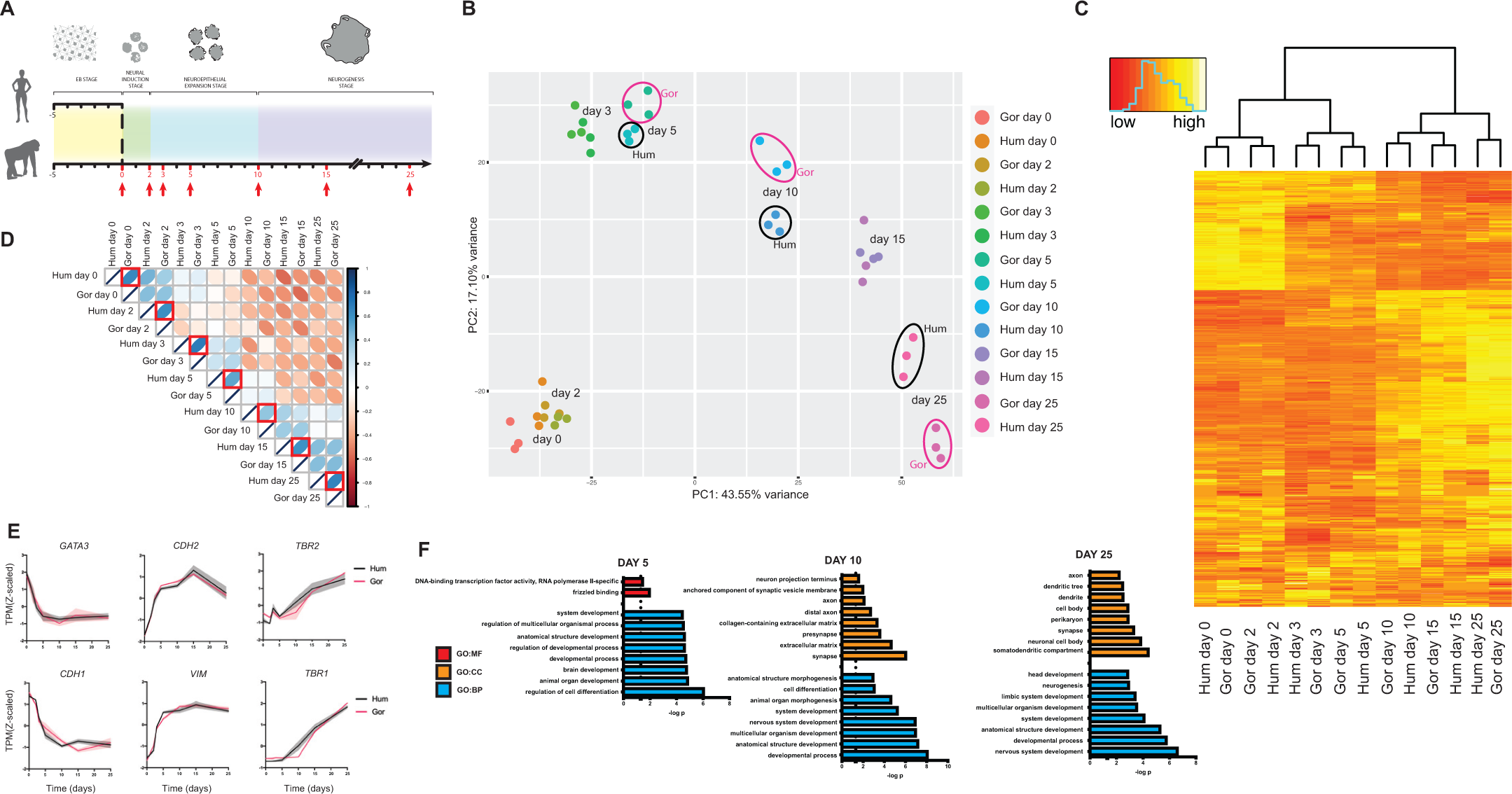
Ape organoids exhibit comparable developmental molecular trajectories. **A.** Schematic of the timeline for human and gorilla brain organoids with RNAseq collection time points shown in red. 3 biological replicates of organoids derived from human (H9) and gorilla (G1) cells were collected at each of the 7 time points. **B.** PCA biplot of PC1 vs PC2 performed on Z-scaled log2-transformed TPMs of the 3000 most variable genes. Samples are color-coded by time point and species. Note samples separating primarily by time point with a slight separation between samples of different species at day 5, 10 and 25, highlighted by ellipses (black ellipses for human and fuchsia for gorilla). **C.** Heatmap with hierarchical clustering based on shared expression pattern (Z-scores of log2-transformed TPMs of the 3000 most variable genes) between samples. The dendogram shows samples clustering by time point. **D.** Pearson’s correlation map using Z-scaled log2-transformed TPMs of all genes. Darker blue depicts stronger positive correlation between samples and darker orange a stronger negative correlation. Red boxes highlight the correlation between species at matched time points. Note slightly lower correlation between species at day 5 and 10. Pearson’s correlation coefficient: r = 0.66 (day 0); 0.63 (day 2); 0.66 (day 3); 0.52 (day 5); 0.40 (day 10); 0.62 (day 15); 0.66 (day 25). **E.** Temporal expression pattern (z-scaled) of characteristic developmental markers show predicted expression dynamics. Non-neural ectoderm markers (*GATA3, CDH1*) are lost rapidly, followed by a gain in neural progenitor markers (*CDH2, VIM*), and a later increase in intermediate progenitor (*TBR2*) and early-born neuron (*TBR1*) markers. Shaded error bar is S.D. **F.** GO term enrichment analysis on the top 300 genes driving species variance at day 5, 10 and 25 (time points separated by PCA in Figure 3B). Shown are the 8 most significant (P<0.05) enrichments for GO categories molecular function (GO:MF), cellular compartment (GO:CC) and biological process (GO:BP).

Principal component analysis of normalized data showed samples of the same time point across species grouped together (Figure 3B). Furthermore, hierarchical clustering on mean z-scores (Figure 3C) and Pearson’s correlation analysis (Figure 3D, S3C) also showed grouping by time rather than species. An initial look at a panel of genes with characteristic developmental roles showed expected patterns of expression in both species with a rapid reduction in the expression of non-neural ectoderm markers (*GATA3, CDH1*) paralleled by a rapid increase in the expression of neural progenitor markers (*CDH2, VIM*), and at later neurogenic stages, an increase in intermediate progenitor (*TBR2*/*EOMES*) and neuronal (*TBR1*) markers (Figure 3E). These data further confirm the proper identity and developmental trajectory of organoids from both species, and demonstrate they are highly comparable.

Although organoids collected at the same time point appeared highly correlated across species, the PCA biplot revealed a degree of separation between species at days 5, 10, and 25 (Figure 3B). To identify what might be contributing to these differences, we performed GO term enrichment analysis on the top 300 genes driving the variance at these time points (Supplementary Data 1). GO terms at all time points were generally associated with nervous system development. However, while the terms associated with the earlier day 5 time point primarily covered organ development and morphogenesis, day 25 GO terms were associated with neuronal compartments such as dendrites and synapses, as well as neurogenesis (Figure 3F). Day 10 seemed to exhibit overlapping GO terms, suggesting these represent an intermediate stage.

Previous studies have reported differences in the rate of neurogenesis and neuronal maturation at later time points in humans relative to other apes (Kanton et al., 2019; Liu et al., 2012; Marchetto et al., 2019). We therefore examined more closely the expression patterns of some of the genes driving the variance at day 25 and saw a steeper rate of increase in genes associated with synaptic formation and maturation in gorilla (Figure S3D). This is consistent with previous studies and suggests that human neurons do indeed mature at a slower rate.

### Species-specific differences in gene expression and biological processes related to cell morphogenesis during NE to RG transition

In addition to a separation in the PCA biplot, the Pearson’s correlation coefficient was also lower at day 5 (r = 0.52) and day 10 (r = 0.40), than any of the other time points (Figure 3D). This weaker correlation further pointed to species-level differences during these stages, which also coincided with our observed differences in cell shape during the transition from NE to RG cells. In order to identify the key players driving these observed cell biological differences, we decided to focus on gene expression changes arising from day 3 to day 5 through day 10, which would cover the time taken in both species for progenitors to transition from NE through tNE to RG morphologies. To assess temporal changes in expression, we analyzed abundantly expressed genes (TPM > 10) with highly reproducible expression patterns between biological replicates for each species (squared difference <6). We also focused on genes with more dynamic expression patterns over the three time points (fold change >1.5 between any two time points in at least one species). This resulted in a list of 2,905 genes that were then subjected to analysis by TCseq (Wu and Gu, 2020), which clustered genes with similar temporal patterns. Focusing on 3 time points meant that temporal patterns could be represented by and separated into 10 clusters with distinct shapes (Figure 4A). During TCseq analysis, biological replicates were kept separate in order to further filter out genes with variable expression patterns, i.e. genes where replicates were split into 3 different clusters in either species. This yielded a list of 2,342 genes per species that were then assigned to the cluster where ≥ 2 of the replicates were located (Supplementary Data 2).

**Figure 4.**
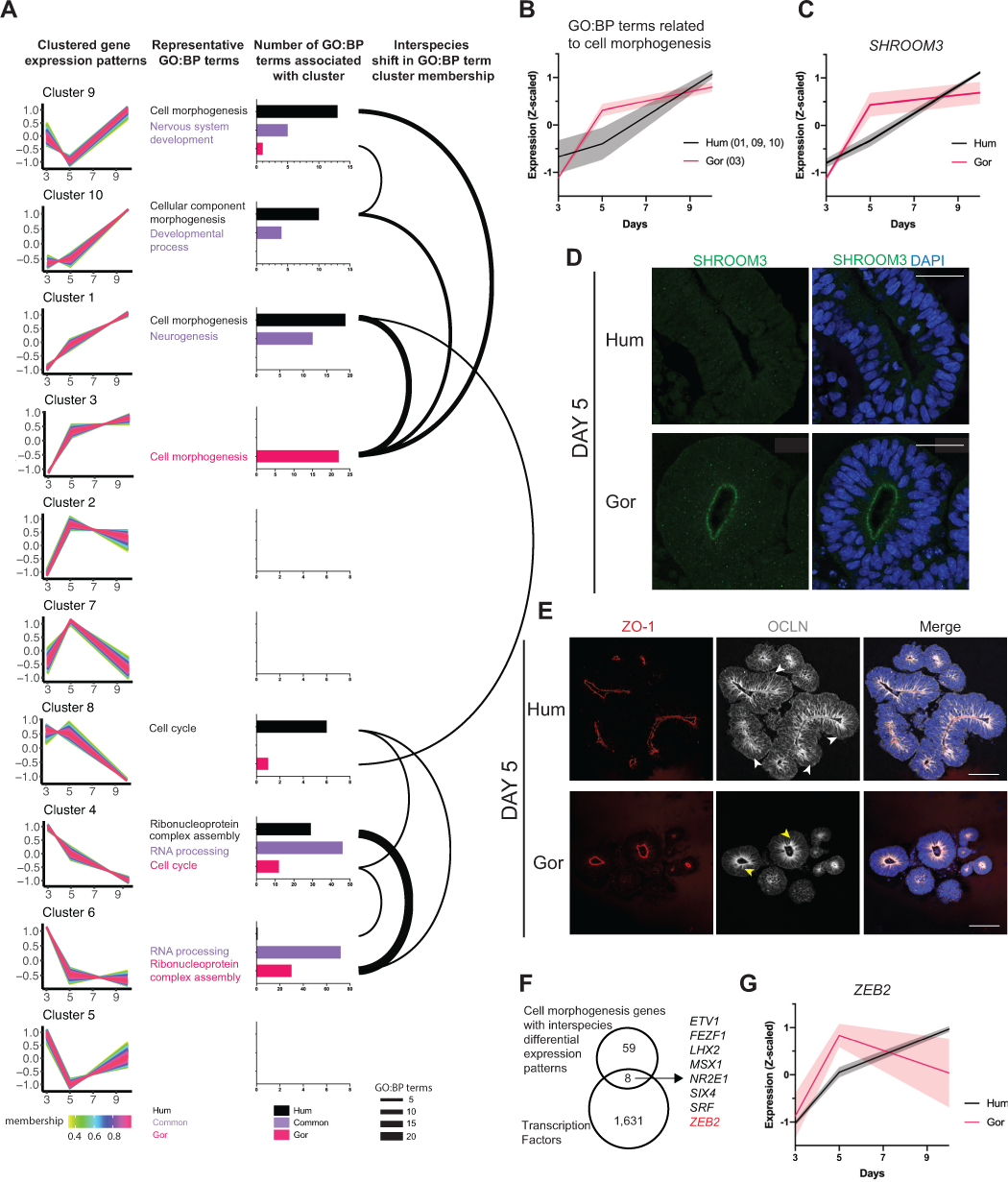
The human neuroepithelium exhibits differential temporal dynamics of morphogenesis genes. **A.** Clustering genes by temporal expression dynamics shows species differences in GO:BP term enrichment. Columns from left to right: Clustered gene expression patterns: TCseq clustering results are visualized, showing genes clustering together based on temporal expression dynamics (z-scaled). The expression pattern of each gene assigned to a cluster is plotted and color-coded by membership value (degree to which data points of a gene belong to the cluster, pink represents high membership values). The 10 clusters are ordered (top to bottom) based on similarity in expression pattern. Representative GO:BP terms: A representative GO:BP term from shared (purple), human-exclusive (black) or gorilla-exclusive (fuchsia) terms for each cluster is shown. Representative terms were not selected for conditions where only one GO:BP term was enriched (i.e. gorilla-exclusive clusters 9 and 8, human-exclusive cluster 6). Number of GO:BP terms associated with cluster: GO term enrichment analysis performed separately on human and gorilla genes present in each cluster identifies biological process (GO:BP) terms enriched (P<0.05) in each cluster. Barplots show the number of enriched GO:BP terms found in both species (purple), exclusively in human (black) or gorilla (fuchsia) per cluster. Axis range: 0 to 8 (cluster 2,5,7,8); 0 to 15 (cluster 9,10); 0 to 20 (cluster 1); 0 to 25 (cluster 3); 0 to 50 (cluster 4); 0 to 80 (cluster 6). Interspecies shift in GO:BP term cluster membership: Weighted arc network graph visualizing interspecies differences in the enrichment/membership of specific GO:BP terms per cluster. The bases of the arc are aligned to both a human (black) and a gorilla (fuchsia) bar from the barplot in the adjacent panel, highlighting the species-specific shifts in expression patterns associated with specific GO:BP terms. Weight/thickness of the arc is dictated by the number of GO:BP terms enriched in a species-exclusive manner “moving” between clusters in the defined pattern. **B.** Mean temporal expression pattern (z-scaled) of genes in clusters enriched for ‘cell morphogenesis’-related GO:BP terms (human clusters 1, 9, 10 and gorilla cluster 3). **C.** Temporal expression pattern (z-scaled) of *SHROOM3*, a gene involved in cell morphogenesis and apical constriction. **D.** Immunofluorescent staining of day 5 organoids for SHROOM3 showing strong apical expression in gorilla (G1) neuroepithelium, but not human (H9) at this time point. Scale bar: 40 μm. **E.** Immunofluorescent staining of day 5 organoids for OCLN showing expression spread along the apicobasal length of human (IMR-90) progenitor cells (white arrowheads) but more limited apically (yellow arrowheads) in gorilla (G1) progenitor cells. DAPI is shown in blue. Scale bar: 100 μm. **F.** Venn diagram summarizing search for cell morphogenesis-related transcription factors with species-specific expression patterns. See methods for more details. **G.** Mean temporal expression pattern (z-scaled) of *ZEB2*, a cell morphogenesis and EMT-related transcription factor, showing peak expression earlier in gorilla (G1) vs human (H9) organoids. Shaded error bars are S.D.

To assign biological functions to the identified temporal patterns, and to see if any of these functions were shifted in human versus gorilla, we performed GO term enrichment analysis on each TCseq cluster, keeping human and gorilla genes separate (Supplementary Data 3). We focused on GO:BP terms found in both species and examined whether each term was found in the same cluster for both species, or different clusters which would indicate a species-specific change in temporal dynamics for that biological function. This revealed a number of GO:BP terms associated with clusters common to both species (Figure 4A), for example GO:BP terms related to “neurogenesis” were found in cluster 1 in both species, with a predictable expression trajectory that gradually increased between day 3 and 10. GO:BP terms linked to “RNA processing”, conversely, were found in clusters of gradually decreasing expression in both species.

There were a number of GO:BP terms associated with a particular expression dynamic in one species but associated with a different pattern in the other species. These represented particularly interesting biological functions as they could be processes governing the differential cell shape changes seen in human versus gorilla. To help visualize the major shifts in temporal expression patterns between human and gorilla, we generated a weighted network graph of GO:BP terms that were uniquely present for one species in a given cluster, but present in a different cluster for the other species (Figure 4A). The largest shift of GO:BP terms was between human cluster 4 and gorilla cluster 6, where terms such as “Ribonucleoprotein complex assembly” and “ncRNA processing” were seen to move from a pattern showing a more gradual decrease in expression in human, to one dropping sharply after day 3 in gorilla (Figure S4A). These terms were linked to the umbrella term, “RNA processing”, which was found in both species in both clusters, but showed higher enrichment in gorilla cluster 6 than cluster 4. Interestingly, we observed a more pronounced drop in the expression of genes related to RNA processing between day 3 and 5 in gorilla compared to human, perhaps pointing to changes in RNA processing such as alternative splicing potentially driving morphological changes. Moreover, several previous studies have highlighted alternative splicing as a key driver of evolutionary differences across primates (Bush et al., 2017).

Another interesting shift in GO terms was “cell cycle” moving from a cluster where expression levels were similar between day 3 and 5 in human (cluster 8), to a cluster where expression levels had already started declining by day 5 in gorilla (cluster 4) (Figure S4B). This suggests that the transition from NE to RG might be coupled with changes in cell cycle length. Previous work in mouse has shown a correlation between progenitor cell differentiation and cell cycle duration, with a progressive lengthening of progenitor cell cycle observed in the developing mouse cortex (Calegari et al., 2005; Takahashi et al., 1995). This also has implications for brain expansion, as a shift to a longer cell cycle would slow down progenitor proliferation.

However, we were most intrigued by the shift in terms such as “cell morphogenesis” and “cellular component morphogenesis”, which were present in cluster 3 of gorilla but in clusters 1, 9 and 10 of human (Figure 4A). Because differences in cell morphogenesis (i.e. cell shape) were precisely what we had observed in our characterization of the organoids from the two species, this shift may help explain the observed differences in tissue size and cell shape. In addition, we found that these terms were associated with genes whose expression increased much more rapidly between day 3 and 5 in gorilla than in human (Figure 4B), matching the more rapid transition in cell shape observed in gorilla at these time points compared with human. In particular, *SHROOM3*, a “cell morphogenesis”-related gene that exhibited differing dynamics in human versus gorilla (Figure 4C), immediately stood out to us due to its known role in inducing apical constriction and elongation, through polarized assembly of both actin and γ-tubulin in various epithelial tissues, including the neuroepithelium (Haigo et al., 2003; Hildebrand and Soriano, 1999; Lee et al., 2007). Immunostaining for SHROOM3 at day 5 confirmed the differences detected by RNA-seq, with strong SHROOM3 localization visible along the whole apical surface in gorilla, whereas apical surfaces of human progenitors showed minimal to no staining (Figure 4D). However, by day 10 strong SHROOM3 staining was seen at the apical surface in both species (Figure S4C).

A closer inspection of the genes contributing to the GO:BP term shift between species, and specifically those displaying earlier onset of expression in gorilla, revealed a group of genes involved in epithelial-to-mesenchymal transition (EMT) (Figure S4D). This was particularly interesting to us as the process of corticogenesis, from NE cells to neurons, is characterized by a progressive and gradual loss of epithelial features, reminiscent of EMT (Singh and Solecki, 2015). One such epithelial feature that is lost is the presence of the tight junction protein OCLN (Aaku-Saraste et al., 1996). Day 3 organoids of both species, when cells are still columnar epithelial in shape, showed OCLN spread along the entire apico-basal axis of individual cells (Figure S4E). By day 10 in both species this OCLN stain was reduced and limited apically, lacking localization along the entire cortical wall (Figure S4F). Day 5 organoids, however, showed species differences in OCLN localization. Human OCLN, similar to day 3, was still expressed along the length of cells. By contrast, gorilla OCLN had already begun transitioning to a more apical localization (Figure 4E), a redistribution that could allow cells to lose their columnar epithelial-like morphologies and exhibit shape changes consistent with the observed tNE. These differences in EMT and cell-cell junction factors could therefore explain the more rapid acquisition of tNE morphologies in ape versus human organoids.

In order to identify upstream regulators that may be responsible for these more global EMT and cell-cell junction differences, we queried the set of genes with robust differential temporal dynamics (Supplementary Data 4) that belonged to the “cell morphogenesis” GO:BP term for genes predicted to be transcription factors. This revealed a set of 8 genes: *ETV1, FEZF1, LHX2, MSX1, NR2E1, SIX4, SRF* and *ZEB2* (Figure 4F). Of these, *ZEB2* stood out in particular due to its well-described role as an EMT master regulator through its repression of epithelial adherens and tight junction genes (Peinado et al., 2007; Stemmler et al., 2019). Furthermore, several comparative genomics analyses have pulled out *ZEB2* as a human gene under selective pressure and exhibiting various human accelerated regions (Erwin et al., 2014; Lindblad-Toh et al., 2011; Pollard et al., 2006) (HARs), which are genomic regions that are highly conserved across mammals, but specifically changed in humans since our divergence from chimpanzees (Pollard et al., 2006). *ZEB2*, which showed a shift from cluster 1 in human to 2 in gorilla, therefore stood out as a potential regulator of cell shape in this context. The expression dynamics of *ZEB2* revealed an earlier peak of expression in gorilla compared with human (Figure 4G), matching the expected expression pattern of a gene that could be initiating the tNE shape transition.

### *ZEB2* is expressed during NE to RG cell transition and this process is delayed in *ZEB2* heterozygous loss-of-function organoids

We decided to focus on *ZEB2* for further functional studies in order to test whether it may be responsible for the species-specific differences in cell shape. First, to confirm the expression differences observed by bulk RNA-seq, we performed an independent expression time-course by quantitative RT-ddPCR on human and gorilla organoids further confirming the shifted RNA expression dynamics of *ZEB2* (Figure 5A) and revealing higher *ZEB2* mRNA levels between day 0 and day 5 in gorilla compared to human. We next performed immunofluorescent staining across various stages of human organoid development to characterize its expression across different cortical cell types over time (Figure 5B-D). ZEB2 was found expressed in dorsal telencephalic progenitors co-expressed with markers such as PAX6 and EMX1 (Fig S5A & B). At day 5, ZEB2 was found in the nuclei of NE cells (Figure 5B). Expression was gradually lost from progenitors as neurogenesis began and by day 20 it started being expressed in newborn DCX+ neurons (Figure 5C). Interestingly, at later stages ZEB2 expression was entirely lost from neurogenic RG cells and remained confined to neurons (Figure 5D). In organoids that were transitioning to neurogenic states, we observed progenitor zones with a salt-and-pepper pattern of ZEB2 expression (Figure 5C). Immunofluorescent staining for markers of committed RG identity, such as BLBP (Figure 5E) and GLAST (Figure S5C) revealed a mutually exclusive pattern of expression between ZEB2 and these markers. These data suggest that ZEB2 expression increases in NE cells during the time window in which they transition to RG cells, but its presence is then lost in neurogenic RG cells at which point it remains exclusively expressed in neurons.

**Figure 5.**
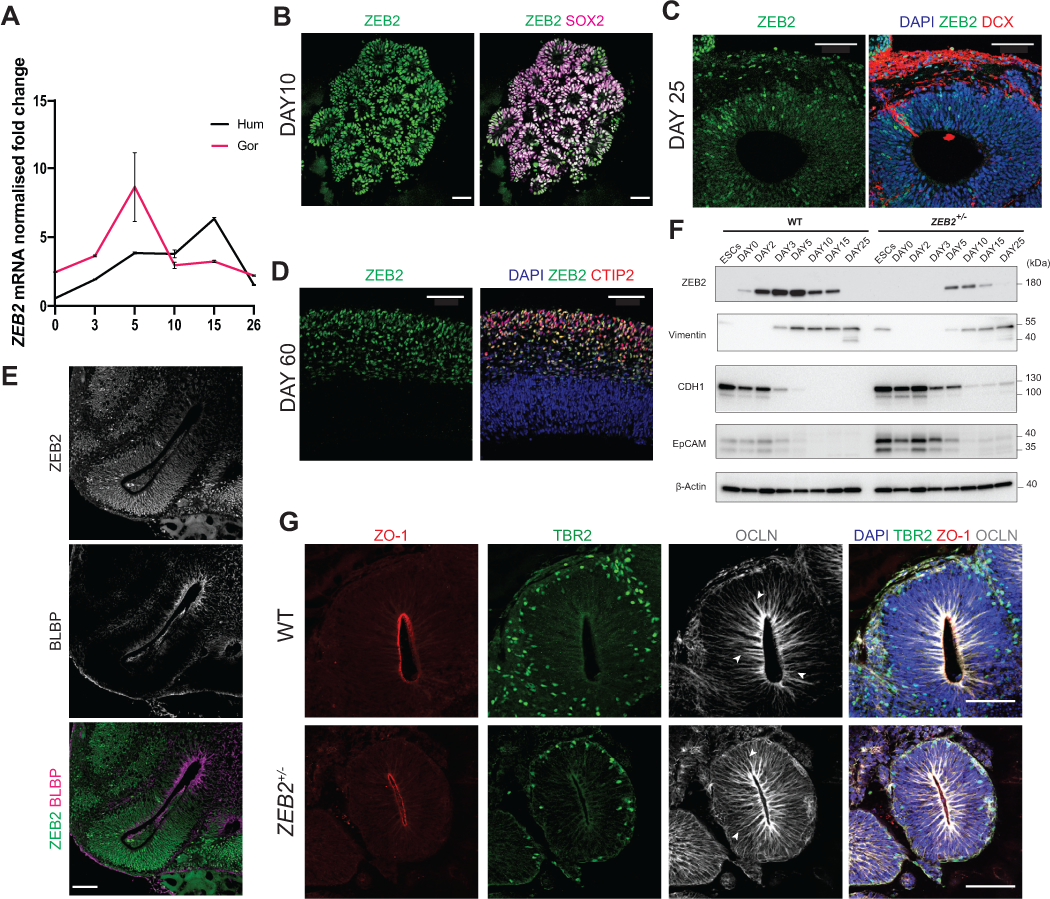
Decreased ZEB2 delays NE transition in human organoids. **A**. RT-ddPCR for *ZEB2* in human (IMR-90) and gorilla (G1) organoids from day 0-25 showing an earlier peak and increased *ZEB2* mRNA levels in gorilla relative to human between day 0-10. The graph reports *ZEB2* mRNA levels normalised to the mRNA levels of EIF2B2 as control (mean±SEM, n=3 technical replicates). **B**. Immunofluorescence image showing ZEB2 expression in SOX2+ progenitor cells in day 10 human (H9) organoid. Scale bar: 50 µm. **C.** Immunofluorescence image showing a salt-and-pepper pattern of ZEB2 expression in the ventricular zone at day 25, after the onset of neurogenesis in human (H9) organoid. DCX (Doublecortin) stains newly born neurons. Scale bar: 100 µm. **D.** Immunofluorescence image of a mature day 60 human (H9) organoid revealing ZEB2 expression in CTIP2+ neurons and absence of ZEB2 staining in the ventricular zone. Scale bar: 100 µm. **E.** Immunofluorescence image of a day 25 human (H9) organoid showing a mutually exclusively pattern of expression between ZEB2 and the committed radial glia marker, BLBP. Scale bar: 100 µm. **F.** Western blot expression time course from PSCs to day 25 organoids of wild-type (WT) and *ZEB2*^+/−^ reveals a decrease in ZEB2 protein levels in *ZEB2*^+/−^ organoids compared to WT across all time points. Loss of ZEB2 protein is accompanied by a delayed onset of expression of the neurogenic radial glial marker Vimentin and a prolonged retention of the epithelial markers CDH1 and EpCAM. **G.** Immunofluorescence images of day 15 WT and *ZEB2*^+/−^ organoids showing increased Occludin immunostaining along the apico-basal axis (arrowheads) of progenitor cells and reduced numbers of TBR2+ cells in *ZEB2*^+/−^ organoids compared to WT. Scale bar: 100 μm.

The pattern of ZEB2 expression pointed to a role in the NE to RG transition. In order to functionally test its requirement, we sought to generate *ZEB2* loss-of-function mutant human ES cells. CRISPR targeting resulted in recovery of only heterozygous cells (Figure S5D-G), suggesting this gene may be necessary for stem cell pluripotency or survival. However, heterozygous loss-of-function mutations in *ZEB2* have been described to cause Mowat-Wilson syndrome, a complex disorder that manifests itself as an array of brain developmental defects with variable penetrance (Mowat et al., 2003). Therefore, we reasoned that heterozygous knockout organoids may be informative, both from an evolutionary point of view, and in understanding this human condition. Cytological and RT-PCR analyses confirmed the cells were healthy and expressed key pluripotency marker genes to the same levels as the parental control (Figure S5H-I), despite showing a reduction in *ZEB2* mRNA levels (Figure S5G). Western blot analysis of day 15 mutant organoids confirmed a reduction in ZEB2 protein compared to control (Figure S5J).

We next tested whether the mutation of *ZEB2* led to changes in tissue architecture consistent with a role in NE to RG transition. Western blot at various time points from day 0 to day 25 revealed a reduced and delayed expression of ZEB2 in mutant organoids compared to control (Figure 5F). This was accompanied by a delay in the loss of the epithelial markers, CDH1 and EpCAM, and delayed expression of the RG and mesenchymal marker, Vimentin (Figure 5F). Furthermore, staining of early neurogenic organoids showed that, while control organoids had redistributed the tight junction epithelial protein OCLN near the apical surface of cells, *ZEB2*+/− mutant organoids retained stronger OCLN staining along the entire apico-basal axis of individual progenitor cells (Figure 5G). These data indicate that mutants preserve epithelial characteristics for a prolonged period. Furthermore, *ZEB2*+/− organoids showed significantly decreased numbers of TBR2+ intermediate progenitor cells compared to wildtype organoids (Figure 5G, S5K), suggesting a potential delay in neurogenesis, which would be consistent with a delayed transition to neurogenic RG.

### Forced *ZEB2* expression in human organoids triggers tNE cell transition leading to a nonhuman ape morphology

The findings so far supported a role for ZEB2 in driving cell adhesion changes underlying the cell shape transition from NE to tNE cells. Next we sought to examine whether this function could explain the species differences seen between human and ape organoids. We first examined ZEB2 and its downstream targets in human and gorilla organoids by western blot, revealing an earlier onset of expression of ZEB2 and Vimentin proteins in gorilla compared to human (Figure 6A). Consistent with a precocious EMT-like transition in gorilla, the epithelial cell adhesion proteins CDH1 and EpCAM showed earlier down-regulation in gorilla compared to human (Figure 6A).

**Figure 6.**
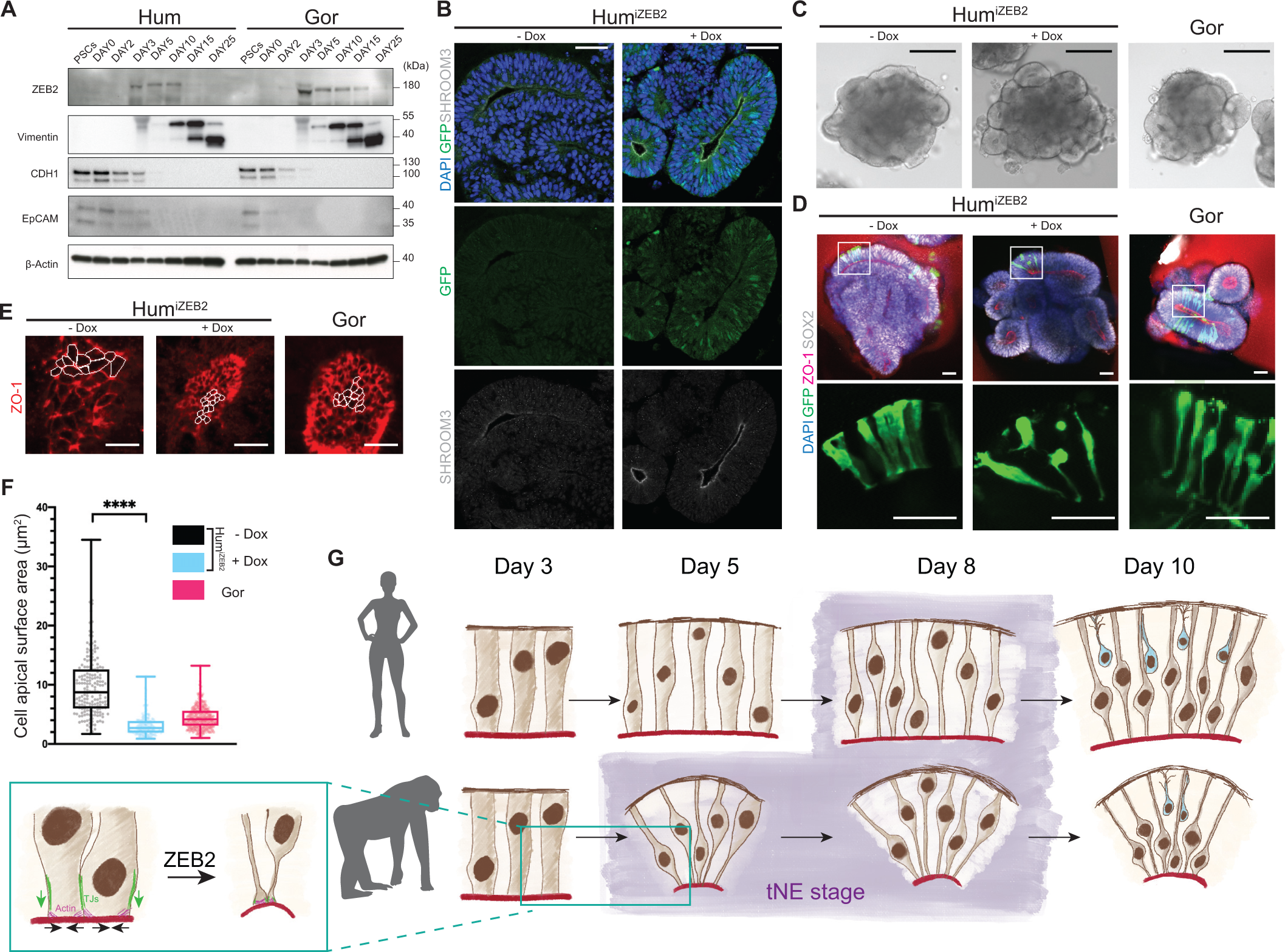
ZEB2 overexpression forces a nonhuman ape phenotype in human organoids. **A.** Western blot expression time course from PSCs to day 25 organoids derived from human (H9) ESCs and gorilla (G1) iPSCs reveals a premature onset and higher levels of ZEB2 protein expression in gorilla compared to human. This is accompanied by a premature expression of the radial glial marker Vimentin, and premature downregulation of the epithelial markers CDH1 and EpCAM in gorilla compared to human. **B.** Immunofluorescent staining of uninduced (-Dox) and induced (+ Dox) Hum^iZEB2^ organoids for GFP and SHROOM3. Note the salt-and-pepper expression of ZEB2-GFP and apical accumulation of SHROOM3 in induced organoids. Scale bar: 50 μm. **C.** Representative brightfield images of day 5 Hum^iZEB2^ and gorilla organoids. Induced Hum^iZEB2^ organoids show smaller neuroepithelial buds that are more rounded in shape, similar to gorilla (G1), while uninduced (-Dox) shows more elongated structures typical of human. Scale bar: 200 μm. **D.** Immunofluorescence images through day 5 whole mount Hum^iZEB2^ uninduced (-Dox), induced (+ Dox) and gorilla (G1) organoids stained for GFP, ZO1 and SOX2. Sparse labelling with viral GFP shows ZEB2 induction triggers the constriction of apicobasal processes in progenitor cells, prematurely adopting a tNE morphology, similar to gorilla progenitor cells at day 5. Scale bar: 50 μm. **E.** Immunofluorescent staining for ZO1 on the surface of apical lumens showing the apical surface areas of individual progenitor cells in day 5 organoids derived from Hum^iZEB2^ uninduced (-Dox), induced (+ Dox) and gorilla (G1) cell lines. Perimeters of some individual progenitor cells of day 5 organoids are delineated in white highlighting the more constricted apical surface areas seen in gorilla and human upon ZEB2 induction compared to uninduced Hum^iZEB2^ organoids. Scale bar: 10 μm. **F.** Quantification of the surface area of individual neural progenitor cells of day 5 organoids show significantly smaller apical surface sizes of induced (+ Dox) vs uninduced (-Dox) Hum^iZEB2^ derived organoids. Measurements were performed on delineated ZO1 cell perimeters such as those seen in Figure 6E. Gorilla measurements from two cell lines (G1, G2) combined are shown for comparison. Mean apical surface area/cell: Hum^iZEB2^ – Dox = 9.62 μm^2^; Hum^iZEB2^ + Dox = 3.08 μm^2^; gorilla (G1,G2) = 4.50 μm^2^. Mann-Whitney U = 2277, **** P<0.0001, two-tailed, n (-Dox) = 180 cells from 8 organoids from 2 independent batches, n (+ Dox) = 199 cells from 8 organoids from 2 independent batches, error bars are min-max values, dots on the boxplot represent individual cells. **G.** Schematic summarizing the morphological changes that occur in human and gorilla derived organoids over time as they transition from NE to tNE to RG. ZEB2 is highlighted as a driver of this progenitor cell differentiation, which involves apical constriction through rearrangements in the actin cytoskeleton and the downregulation of epithelial features, notably tight-junction proteins (TJs).

Given these species differences in ZEB2 protein expression, we next tested whether changing *ZEB2* expression dynamics to match that of the gorilla would be sufficient to trigger an earlier transition to tNE cells in human organoids. We generated a human cell line for doxycycline inducible overexpression of *ZEB2-GFP* (Figure S6A-B). We observed robust and tight expression upon induction, both by western blot and immunofluorescent staining (Figure S6C-D). The nuclear localization of ZEB2-GFP in doxycycline treated stem cells suggested that C-terminal tagging did not interfere with the normal intracellular sorting of the protein (Figure S6D). Following doxycycline treatment in stem cells we could observe a reduction in EpCAM and CDH1 staining accompanied by an increase in VIM and CDH2 staining (Figure S6E-F) relative to the un-induced control.

We next generated organoids from these inducible *ZEB2* human (Hum^iZEB2^) cells, and treated with doxycycline to induce *ZEB2* expression at an earlier stage, as seen in gorilla. This induction resulted in a salt-and-pepper expression of ZEB2-GFP in the nuclei of NE cells by day 5 (Figure 6B). Examination of tissue architecture at day 3 revealed indistinguishable morphology between uninduced Hum^iZEB2^, induced Hum^iZEB2^, and gorilla organoids (Figure S6G). By day 5, however, neuroepithelial buds generated in induced Hum^iZEB2^ organoids appeared more similar to gorilla in shape, appearing smaller and more rounded, while uninduced Hum^iZEB2^ organoids exhibited buds with more elongated shapes typical of human organoids (Figure 6C, Figure S6G). This change in morphology was also accompanied by a change in expression and localization of SHROOM3, the actin regulator identified to exhibit precocious expression in gorilla. Whilst uninduced Hum^iZEB2^ cells exhibited only faint staining for SHROOM3, induced Hum^iZEB2^ cells exhibited SHROOM3 accumulation at the apical surface (Figure 6B), as seen in day 5 gorilla organoids (Figure 4D).

Finally, we performed sparse viral labeling with GFP revealing that SOX2+ NE cells of induced Hum^iZEB2^ organoids were less columnar and more tNE-like in shape than un-uninduced cells, with a thinning of apical and basal processes (Figure 6D). This morphology was highly similar to gorilla tNE cells in day 5 organoids (Figure 6D). We quantified the changes in cell shape and apical constriction by measuring apical surface area using ZO1 to outline the apical periphery (Figure 6E). This revealed a remarkably similar apical constriction in human *ZEB2* overexpressing organoids to gorilla organoids, which was 1/3 the size of the apical domain of NE cells in un-induced human organoids (Figure 6F). These findings suggest that premature expression of *ZEB2* in human organoids is sufficient to recapitulate the precocious tNE cell shape change seen in gorilla organoids (Figure 6G), thus revealing ZEB2 as a key regulator of species-specific human neuroepithelial expansion.

## Discussion

These results provide novel insights into how NE cells transition into neurogenic RG cells, highlighting an intermediate tNE cell stage, hitherto unidentified in the mammalian neocortex. Interestingly, a transitioning NE cell type has been described in Drosophila optic primordia during the differentiation of NE to neuroblast cells (Orihara-Ono et al., 2011). This transition involves features analogous to those we have observed in human and ape organoids, displaying changes in morphology, apical constriction and a loss of epithelial characteristics, raising the intriguing possibility that such a transition may be evolutionarily conserved. Furthermore, the temporal progression of events involved in this transition point to a shift in cell morphology preceding other changes associated with RG identity. RNAseq further supports this conclusion, revealing the acquisition of RG molecular identity (expression of glial markers) and cell fate (generation of neurons) after the shift in morphology. This suggests that cell shape may be a key initiator in this developmental transition.

The protracted shift to the tNE morphotype in humans is particularly intriguing with regard to human brain evolution, as it may explain to a large degree the linear scaling of the human brain compared with apes. An evolutionary modification at such an early developmental stage would be expected to affect all subsequent steps of neurogenesis, and lead to a proportional increase in all cell types. Furthermore, cytoarchitecturally, this would be expected to lead not to a thickened cortical grey matter, but rather to a more expanded brain in the tangential dimension, which is precisely what is seen in comparative neuroanatomy to other apes. Previous observations of human cerebral organoids compared with chimpanzee have focused on later stages but similarly observed protracted human development (Kanton et al., 2019), pointing to a common thread in human evolution: timing. These studies are therefore complementary, but the effect on brain size and cellular make-up differ, and the mechanism of these later stage differences remains to be identified.

Functional studies are necessary in delineating mechanism. We therefore took our observations on developmental timing a step further, and performed functional studies of a key factor we identified by RNAseq, *ZEB2. ZEB2* was previously shown to be involved in early germ layer fate (Chng et al., 2010; Eisaki et al., 2000; Stryjewska et al., 2017), neural tube morphogenesis (Miyoshi et al., 2006), neural crest differentiation (Yasumi et al., 2016), and cortical laminar fate determination (Epifanova et al., 2019), but it is most well-known as a central node of the epithelial to mesenchymal transition (EMT) control network (Fardi et al., 2019). In many respects, the transition from NE to tNE to RG resembles a partial EMT process with a progressive change in cell morphology, apical constriction and a reduction in cell-cell junctions. Indeed, even in more classic EMT scenarios, such as cancer metastasis, there is increasing recognition that EMT is not a binary switch, but rather that there are several intermediate stages (Pastushenko and Blanpain, 2019). Additionally, several studies to date have shown that EMT is linked to cytoskeletal changes (Morris and Machesky, 2015) and our observations of increasing expression and accumulation of SHROOM3 apically demonstrates that NE transition is coupled to such cytoskeletal rearrangements (Chu et al., 2013). Our findings also support a link between ZEB2 and cytoskeletal targets, and although the specific nature of this link remains to be elucidated, several reports have shown cross-regulation between ZEB2 and the actin cytoskeleton (Wiles et al., 2013; Yuan et al., 2019).

In conclusion, we have taken an intriguing observation involving timing of a novel cellular morphotype in early human brain development, worked out the key steps in this transition in human compared with nonhuman ape, and identified a crucial regulator of this process. Importantly, the *ZEB2* locus exhibits signatures of evolutionary selection, and recent scATAC-seq data has identified 6 differentially accessible regions between human and chimp associated with *ZEB2* (Kanton et al., 2019). These changes in noncoding regions point to differences in gene regulation, which would be consistent with the expression level differences we observe between human and ape organoids. Finally, we were able to mimic such expression level differences and effectively “gorillize” the human organoids, strongly pointing to *ZEB2* as a key regulator of human brain evolution. Overall, our findings suggest that a relatively simple evolutionary change in a regulator of cell shape can have major consequences in brain evolution.

## Materials and Methods

### Cell lines

One human ESC line (H9), one human iPSC line (iPS(IMR90)-4, shortened to IMR-90 in this study), one chimpanzee iPSC line (C3651, shortened to Chmp) and two gorilla iPSC lines (goiPSC clone 1 and gorC1, shortened to G1 and G2 respectively) were used in this study. All cell lines were female. H9 and iPS(IMR90)-4 were purchased from WiCell. C3651 was a gift from Yoav Gilad (Gallego Romero et al., 2015). goiPSC clone 1 was previously published (Wunderlich et al., 2014). We generated gorC1 using peripheral blood mononuclear cells (PBMCs) isolated from leftover blood collected from a female western lowland gorilla during a routine check-up at Twycross zoo. Briefly, PBMCs were isolated using Lymphoprep (Stem Cell Technologies, 07801), followed by expansion for 9 days in Expansion Media composed of Stemspan H3000 (Stem Cell Technologies, 09850) supplemented with 50μg/ml ascorbic acid (Sigma, A4403), 50 ng/ml SCF (Miltenyi, 130-096-692), 10 ng/ml IL-3 (Invitrogen, PHC0035), 2 U/ml EPO (R&D Systems, 287-TC-500), 40 ng/ml IGF-1 (Miltenyi, 130-093-885), 1 μM Dexamethasone (Sigma, D8893-1MG). On day 10, expanded PBMCs were transduced with non-integrating Sendai reprogramming virus (Cytotune 2.0, Thermo Fisher, A16517) according to manufacturer’s instructions added to the Expansion Media. The next day, the media was changed and two days later the cells were collected and plated on gamma irradiated MEF feeders (Gibco, A34181) in human ES media (DMEM/F12 (Invitrogen, 11330-032) with 20% knockout serum replacement (Invitrogen, 10828-028), 1:100 Glutamax (Invitrogen 35050-038), 1:100 NEAA (Sigma, M7145), 3.5μl/500ml 2-mercaptoethanol, 20 ng/ml bFGF (Peprotech, 100-18B)) supplemented with ascorbic acid, SCF, IL-3, EPO, IGF-1, and Dexamethasone at the above concentrations along with iPS Boost Supplements II (Millipore, SCM094). Two days later (5 days after induction), media was changed to human ES media without PBMC supplements but with Boost Supplements. Cells were maintained in hES + Boost until colonies began to form (10 days after induction). The small colonies were then split off MEFs 15 days after induction using collagenase and a cell lifter and replated on Matrigel (Corning) coated dishes in StemFlex (Thermo Fisher, A334901) for further selection and propogation. Human ESCs used in this project were approved for use in this project by the U.K. Stem Cell Bank Steering Committee and iPSCs and ESCs were approved by an ERC ethics committee and are registered on the Human Pluripotent Stem Cell Registry (hpscreg.eu). Nonhuman primate cells were approved by the Animal Welfare and Ethical Review Body (AWERB) of the MRC-LMB. RT-PCR (refer to the *‘*Digital droplet PCR (ddPCR) and RT-ddPCR’ section of the methods) confirmed pluripotent gene expression (Figure S1F). All cell lines were maintained in StemFlex on Matrigel coated plates. Cells were passaged every 3-4 days using 0.7 mM EDTA. HEK293 cells were a gift from Dr Harvey McMahon and were cultured in high-glucose DMEM supplemented with GlutaMAX (Thermo Fisher, 31966047) and 10% FBS (Thermo Fisher, 10270106) and passaged once a week using TrypLE (Thermo Fisher, 12563029).

### Plasmid Constructs

All oligos used for cloning are listed in Supplementary Table 1. All PCR amplification steps were done using Q5 polymerase (NEB, M049L) and TOP10 chemically competent *E. coli* (Thermo Fisher, C404010) were used for transformation. In order to generate the construct pT2-CAG-fl-STOP-fl-fGFP, the fl-STOP-fl cassette was excised from the pHB4 vector (a gift from Dr Jason Chin) by restriction digestion with EcoRI and AscI and ligated into the pT2-CAG-fGFP (Addgene plasmid #108714) opened with the same restriction enzymes. The construct AAVS1-Puro-CAG-fl-STOP-fl-Cas9 was generated as follows; the backbone was amplified from the Puro-Cas9 donor plasmid (a gift from Dr Danwei Huangfu, Addgene plasmid #58409)(González et al., 2014) with primers AAVS1_CAG_fl_STOP_fl_F & R to introduce MluI (NEB, R3198S) and KpnI (NEB, R3142S) restriction sites and blunt-end ligated to the CAG-fl-STOP-fl cassette, excised from the construct pT2-CAG-fl-STOP-fl-fGFP with restriction enzymes AflI (NEB, R0520S) and AscI (NEB, R0558S) and blunted with the Quick Blunting™ Kit (NEB, E1201). The resulting intermediate plasmid was then digested with MluI and KpnI and ligated to the cassette containing Cas9 and the bGH-poly(A) sequence amplified from pAAV-Neo_CAG-Cas9 (a gift from Dr Ludovic Vallier, Addgene plasmid # 86698)(Bertero et al., 2016) with primers Cas9_β_globin_pA_F & R. All ligations to generate this construct were done with the Quick Ligation™ Kit (NEB, M2200S) and the final construct was sequence validated. The construct AAVS1-Neo-TRE-CMV-CRE-rtTA was generated as follows; the tight TRE promoter was amplified from the Puro-Cas9 donor plasmid (Addgene plasmid #58409) with primers TRE_F &R and inserted into the EcoRV restriction site of AAVS1-Neo-M2rtTA (a gift from Dr Rudolf Jaenisch, Addgene plasmid # 60843)(DeKelver et al., 2010) to produce the intermediate construct AAVS1-Neo-M2rtTA-TRE. A fragment containing CMV-2TO-MCS-bGH-poly(A) was PCR amplified from pcDNA4/TO (Thermo Fisher, V102020) using primers CMV_bGH_F and _R, and inserted into the EcoRV restriction site of AAVS1-Neo-M2rtTA. The fragment comprised between the SalI and AleI restriction sites was replaced with the Cre recombinase cDNA amplified from pCAG-Cre (a gift from Dr Connie Cepko, Addgene plasmid # 13775)(Matsuda and Cepko, 2007) using primers Cre_SalI_F and Cre_KpnI_R. From the resulting plasmid a fragment containing CRE-bGH-poly(A) was excised with restriction enzymes SalI (NEB, R3138S) and AleI (NEB, R0634) and inserted into AAVS1-Neo-M2rtTA-TRE digested with the same restriction enzymes. All ligations to generate this construct were done with the Quick Ligation™ Kit (NEB, M2200S). The final construct was sequence validated. The PX458-AAVS1 construct used to target the AAVS1 locus was generated by cloning the AAVS1_sgRNA sequence into pSpCas9(BB)-2A-GFP (PX458) (a gift from Dr Fang Zhang, Addgene plasmid # 48138) as previously described by Ran et al. (2013). In order to generate the construct for inducible *ZEB2* expression from the AAVS1 site the coding sequence of the human *ZEB2* transcript variant 1 (NM_014795.3) was purchased as an ultimate ORF clone in the pENTRY221 (Thermo Scientific, #IOH53645). The *ZEB2* ORF was amplified with primers ZEB2_IndOE_F & _R and tagged with GFP-Flag by Gibson assembly into the pT2-CAG-MCS-2A-Puro plasmid linearized by restriction digestion with EcoRI (NEB, R3101S) and AgeI (NEB, R3552S). The GFP fragment was amplified and C-terminally Flag-tagged using primers GFP_IndOE_F & R and the construct pT2-CAG-fGFP (Addgene plasmid #108714) as template. While the original design aimed to fuse ZEB2 to GFP via a 4x(GGGGS)-linker motif, Gibson assembly resulted in motif duplication and produced a 6x(GGGGS)-linker between the two coding sequences. The resulting plasmid, pT2-CAG-ZEB2-GFP-Flag-2A-Puro, was used as template in a PCR reaction with primers ZEB2_GFP_Flag_IndOE_F & _R. The resulting fragment encoding ZEB2-GFP-Flag was then digested with restriction enzymes AgeI (NEB, R3552S) and KpnI (NEB, R3142S) and ligated into the CAGs-MCS-Enhanced Episomal Vector (EEV) (System Biosciences, EEV600A-1) linearized with AgeI and KpnI, using T4 ligase. Next, using primers ZEB2_AAVS1 IndOE_F &_R a fragment containing ZEB2-GFP-Flag-WPRE-poly(A) and overhangs containing MluI and FseI restriction sites at the 5’ and 3’ end, respectively, was generated. The construct AAVS-Puro-CAG-fl-STOP-fl-Cas9 was linearized with MluI (NEB, R3198S) and FseI (NEB, R0588S) to remove the Cas9 coding sequence, which was then replaced with the fragment encoding ZEB2-GFP-Flag-WPRE-poly(A) by ligation with T4 ligase. The *ZEB2* ORF was verified after every cloning step by sequencing using primers ZEB2_Seq_1-9. For simplicity, this overexpressing construct is referred to as ZEB2-GFP throughout the text. For CRISPR-Cas9 knockout of *ZEB2* in Human and Gorilla PSCs guide RNAs were designed using the online tool at http://tools.genome-engineering.org and the sequences are listed in Supplementary Table 1 as ZEB2*_*sgRNA_1 & _2. Cloning of the guides was performed as outlined by Ran et al. (2013); briefly, the sense and antisense strand oligos for ZEB2_sgRNA_1 & 2 were annealed and phosphorylated, and the duplexes were cloned into pSpCas9n (BB) (PX460) and pSpCas9(BB)-2A-GFP (PX458) (a gift from Dr Fang Zhang, Addgene plasmid #48873 and #48138, respectively) by BbsI (NEB,R3539S) digestion and ligation with T4 ligase. Colonies were sequence validated using the U6_F primer.

### Transgenic cell lines

For establishment of the H9 *ZEB2*^*+/−*^ line, plasmids pSpCas9n(BB)-*ZEB2*-Guide-A &-B (0.5 µg of each plasmid) were electroporated into 1×10^6^ H9 cells using the Human Stem Cell Nucleofector Kit 1 (Lonza, VPH-5012). Following electroporation, cells were grown in one well of a 24-well plate, reduced to a single-cell suspension and seeded into a 96-well plate at a density ranging between 1000 – 20 c/w in mTesR^TM^1 supplemented with 1 nM ROCK inhibitor (BD Biosciences, 562822). Alternatively, cells were seeded at a density of 0.5 c/w in StemFlex supplemented with RevitaCell supplement (Thermo Fisher, A2644501). Once the cells reached ∼80% confluence the 96-well plate was split to two replica plates, one used for screening by ddPCR and the other used for further expansion. Mutant screening relied on a droplet digital PCR (ddPCR) drop-off assay (Findlay et al., 2016). Screening was performed as described in the *‘*Digital droplet PCR (ddPCR) and RT-ddPCR’ section of the methods. The mutant colonies were characterized by sequencing following cloning of the edit-sites into the pJET1.2 vector (using primers ZEB2_Cas9screening_F & _R), PCR followed by DNA-PAGE (using primers ZEB2_DNAPAGE_ F & _R), karyotype analyses (outsourced to Cell Guidance Systems) and RT-PCR of pluripotency markers (details in the ‘PCR analysis’ section of the methods).

For establishment of the H9 doxycycline-inducible ZEB2 line (shortened to Hum^iZEB2^), plasmids AAVS1-Neo-TRE-CMV-Cre-rtTA (2 µg), AAVS1-Puro-CAG-fl-STOP-fl-ZEB2-GFP-Flag-WPRE-poly(A) (2 µg) and PX458-AAVS1 (3 µg) were electroporated into 1.3×10^6^ H9 cells using the Human Stem Cell Nucleofector Kit 1 (Lonza, VPH-5012). Following electroporation the cells were seeded across 3-wells of a 6 well plate coated with Matrigel in StemFlex supplemented with RevitaCell supplement (Thermo Fisher, A2644501). After 3 days from electroporation the cells were selected with StemFlex supplemented with G418 (25 µg/ml) and Puromycin (0.5 µg/ml). After approximately two weeks in selection medium, single colonies were picked and screened using three primers pair combinations; AAVS1_F & _R (AMP_1 in Figure S6), AAVS1_F & Puro_R (AMP_2 in Figure S6), AAVS1_F & Neo_R (AMP_3 in Figure S6), listed in Supplementary Table 1. Individual colonies were tested for ZEB2-GFP transgene induction by western blot and immunofluorescence following treatment with StemFlex supplemented with 1.7 µg/ml Doxycyline for 6 days.

### Cerebral organoid generation

Cerebral organoids with telencephalic identity from human and ape cells were generated as previously (Lancaster et al. 2017) with the following modification: Aggrewell 800 plates (StemCell Technologies) containing 300 microwells, each 800 μm in size, per well were used to reproducibly generate organoids in multiples of 300. Following previously described directions for EB generation in Aggrewell plates (StemCell Technologies Document #DX21397), 6 × 10^5^ cells in 1.5 ml media were plated per well, i.e. 2,000 cells per EB. STEMdiff Cerebral Organoid Kit (StemCell Technologies, 08570) was used for organoid culture, with the addition of 1:1000 Fungizone and 1:100 Penicillin-streptomycin to avoid contamination risk associated with repeated plate usage. The timing of the protocol (StemCell Technologies Document #DX21849) was followed, with the exception of chimpanzee embryoid bodies being moved to neural induction media 2 days earlier than described. Organoids were fed every 2 days during the Aggrewell period prior to Matrigel embedding, using an electronic pipette set to the slowest speed in order to avoid flushing EBs out of microwells. Matrigel embedding was based on a previously described method (Qian et al., 2018) with minor modifications. Briefly, organoids were removed from their well, resuspended in 300 μl Expansion media and split into 3 tubes where 150 μl Matrigel was added resulting in a 3:2 Matrigel dilution. The contents of each tube were then spread into a well of an ultra-low attachment 24-well plate (StarLab, CC7672-7524). After a 30-minute incubation at 37 °C, 1 ml expansion media was added per well. Experiments on *ZEB2*^*+/−*^ organoids were generated according to the STEMdiff Cerebral Organoid Kit (StemCell Technologies, 08570) or alternatively with fibrous microscaffolds and media formulation as previously described (Lancaster et al., 2017). To achieve sparse labelling of neural progenitor cells, 5 μl CytoTune emGFP Sendai fluorescence reporter (Thermo Fisher, A16519) was added to Aggrewells when the organoids were switched to neural induction medium. For ZEB2 overexpression, organoids generated from the H9 doxycycline-inducible ZEB2 (Hum^iZEB2^) cell line were treated with 1.7 μg/ml doxycycline (Merck, PHR1145) during the first 5 days of the protocol.

### Histological and immunohistochemical analysis

Organoids were fixed in 4% PFA either overnight at 4 °C or at room temperature for 20-60 min, and washed in PBS (3×10 min). Samples for cryostat processing were incubated overnight in 30% sucrose in 0.2 M PB (21.8 g/l Na2HPO4, 6.4 g/l NaH2PO4 in dH2O), embedded in gelatin (7.5% gelatin, 10% sucrose in 0.2 M PB), plunge frozen in 2-methylbutane (Sigma-Aldrich, M32631) at ∼-40 °C and sectioned at a thickness of 20 μm. Both sectioned and whole mount samples were stained as previously described (Lancaster et al., 2013).

### Imaging and image analysis

For live imaging, organoids were placed on an imaging dish (2BScientific, 6160-30) and were held in place by a drop of Matrigel that was allowed to polymerize before addition of expansion media (StemCell Technologies, 08570). Time-lapse movies were acquired on an Andor Revolution Spinning Disk microscope in an incubation chamber set to 37 °C and 5% CO_2_. Images were taken at 4% laser power every 20 minutes and stacked at 2 μm intervals. All confocal imaging on fixed samples was done on a Zeiss 780 or 880 confocal microscope. Z-stacks of whole mount organoids were generated by taking images at 8 μm intervals. To calculate the apical luminal surface area of organoids, folders containing the Z-stacks of ZO1 stained organoids as an image stack were carried over to Matlab (Mathworks). The largest lumen per organoids was measured using a tailor-made code that delineates the perimeter of the lumen in each Z-stack by detecting fluorescent intensity and then combines the delineated perimeters into a 3D structure from which the surface area is calculated. This code is publicly available (https://github.com/esriis/organoid-lumen-segmentation). The apical surface area of individual cells was measured from sections of organoids where cells were stained positive for SOX2 and sectioned at their apical surface. ZO1 stain was used as a marker of cell perimeters and FIJI (Schindelin et al., 2012) was used to manually delineate perimeters and perform surface area measurements. TBR2 positive cells (Figure S5K) were counted using a FIJI plug-in generated by Johannes Schindelin.

### Immunoblotting

Cell and organoid samples were washed twice in ice-cold PBS, pelleted by centrifugation (500g, 3 min) and lysed with modified-RIPA (mRIPA: 1% Triton-X, 0.1% SDS, 150 mM NaCl, 50 mM Tris pH 7.4, 2 mM EDTA, 12 mM sodium deoxycholate) supplemented immediately prior to lysis with protease (Thermo Fisher, 78430) and phosphatase (Sigma-Aldrich, 4906845001) inhibitors. The protein concentration of the samples was measured using the Quick Start Bradford Dye Reagent (Bio-Rad, 5000205). Between 3-20 µg of total protein per sample were resolved by SDS-PAGE (4-20% gels) and transferred to Amersham Hybond P 0.45 PVDF blotting membranes (GE Healthcare, 10600023). Membranes were blocked overnight at 4 °C in 5% milk or 5% BSA in PBST. Specific blocking conditions were optimized for each antibody during the initial validation stages. Primary antibodies were incubated overnight at 4 °C. HRP-linked goat anti-rabbit (Dako, P0448, 1:3000) and rabbit anti-mouse (Dako. P0161, 1:3000) secondary antibodies were incubated for ∼1 hr at room temperature. The blots were developed using ECL Prime enhanced chemoluminescent detection reagent (GE Healthcare, RPN2232) or alternatively SuperSignal™ West Femto chemoluminescent substrate (Thermo Fisher, 34094) and X-ray films (Photon Imaging Systems Ltd, FM024) and developer or a Gel Doc XR^+^ system.

### Antibodies

Primary antibodies used for protein detection, with their corresponding dilutions for immunofluorescence (IF), western blotting (WB) and WB blocking conditions were as follows: mouse anti-β-actin (Abcam, 8226, WB 1:2000 in BSA), mouse anti-ZEB2 (Origene, TA802113, IF 1:150, WB 1:2000 in milk), sheep anti-TBR2 (R&D Systems, AF6166, IF 1:200), mouse anti-CDH2 (BD Biosciences, 610920, IF 1:500, WB 1:1000 in milk), mouse anti-CDH1 (BD Biosciences, 610181, IF 1:500, 1:1000 in milk), rabbit anti-Occludin (Abcam, ab31721, IF 1:200, WB 1:1000 in milk), rabbit anti-EMX1 (ATLAS Antibodies, HPA006421, IF 1:100), rabbit anti-EMX1 (Origene, TA325087, WB 1:1000 in BSA), rabbit anti-BLBP (Abcam, ab32423, IF 1:200), rabbit anti-GLAST (Abcam, ab416, IF 1:200), goat anti-DCX (N-19) (Santa Cruz, sc-8067, IF 1:300), mouse anti-ZO1 (BD Biosciences, 610966, IF 1:300), chicken anti-GFP (Thermo Scientific, A10262, IF 1:500), rabbit anti-GFP (Abcam, ab290, WB 1:1000 in milk), rabbit anti-EpCAM (Abcam, ab71916, IF 1:300, WB 1:1000 in milk), mouse anti-Vimentin (V9) (Santa Cruz, sc-6260, IF 1:200, WB 1:1000 in BSA) rabbit anti-PAX6 (Abcam, ab195045, 1:200), rabbit anti-SOX2 (Abcam, ab97959, IF 1:200), rabbit anti-SHROOM3 (ATLAS Antibodies, HPA047784, IF 1:200). Alexafluor 405, 488, 568 and 647 secondary antibodies (Thermo Fisher) were used for detection of primary antibodies in IF.

### PCR analysis

For applications aimed at amplicon size comparison GoTaq Green Master Mix (Promega, 9PIM712) was used according to the manufacturer’s guidelines. For molecular cloning or any other application requiring high sequence fidelity Q5 High Fidelity 2X Master Mix (NEB, M0492S) was used. PCR analysis of pluripotency markers was done using the Human Pluripotent Stem Cell Assessment Primer Pair Panel (R&D Systems, SC012) and amplification was done using GoTaq Green. Novex TBE 10% gels (Thermo Fisher, EC6275BOX) were used for DNA-PAGE analysis of *ZEB2*^+/−^ mutant gDNA. The primers used were ZEB2_DNAPAGE_F & R (Supplementary Table 1), GoTaq green was used for amplification and 1X SYBR Gold Nuclei Acid Stain (Thermo Scientific, S11494) was used for detection. Samples were prepared as outlined in the PAGE gels technical sheet.

### Digital droplet PCR (ddPCR) and RT-ddPCR

In order to detect CRISPR-mutants and perform enrichment by sib-selection a TaqMan-based ddPCR drop-off assay was designed (Findlay et al., 2016). An amplicon of 198 bp overlapping the edited genomic region (GRCh38/hg38 chr2:144,517,275-144,517,351) was produced using primers ZEB2_Cas9screening_F & R. The assay was designed so as to have a 5’-HEX-labelled 3’-BHQ1 probe binding to the edited site (i.e. ZEB2_drop-off_probe) and a 5’-FAM-labelled 3’-BHQ1 probe binding to both edited and WT amplicons (i.e. ZEB2_reference_probe). The specific sequence of primers and probes used in the assay are listed in Supplementary Table 1. The assay was performed using the ddPCR Supermix for Probes (Bio-Rad, 1863024) as described in the product’s technical bulletin. Briefly, reaction mixes (20 µl/reaction) were prepared as follows: 100 nM primers, 200 nM probes, 10 U MseI (NEB, R0525S), 1x ddPCR supermix for probes and 50-300 ng of genomic DNA (gDNA). The reactions were loaded into DG8 cartridges (Bio-Rad, 1864008) with droplet generation oil for probes (Bio-Rad, 1863005), the cartridge was then fitted with the DG8 gaskets (Bio-Rad, 1863009) and run on the QX200 droplet generator (Bio-Rad, 10031907). The droplet-oil emulsions were transferred to a ddPCR-compatible 96-well plate that was sealed with the PX1 PCR Plate Sealer (Bio-Rad) and the PCR reaction was run on a C1000 touch thermal cycler following the PCR protocol detailed in the technical bulletin. After thermal cycling, data were acquired on the QX200 Droplet Reader (Bio-Rad) using the QuantaSoft Software (Bio-Rad). Negative and positive control samples for the assay were gDNA extracts from HEK293 cells and HEK293 cells transfected with plasmids pSpCas9(BB)-2A-GFP-*ZEB2-*guide-A & -B, respectively. RT-ddPCR was used in order to quantify target gene expression levels. Briefly, RNA was isolated from organoid tissue using TRIZOL reagent (Thermo Fisher, 15596026). cDNA was synthesized from 240 ng-1 µg total RNA using the SuperScript™ III first-strand synthesis supermix (Thermo Fisher, 18080400) following the product’s manual. Primers used for RT-ddPCR on *ZEB2* and *EIF2B2* are detailed in Supplementary Table 1 and were manually designed to bind across gorilla and human. Amplicons were validated and thermocycling conditions were optimized for individual targets. After reverse transcription, cDNA concentration was not measured – for samples synthesized starting from 240 ng of RNA 0.33 µl of cDNA were used per reaction, while samples synthesized starting from 1 µg of RNA were first diluted 1:5 and 0.33 µl were used per reaction. Detection relied on EvaGreen chemistry (Bio-Rad, 1864034) and because the loading control (i.e. *EIF2B2*) was run separately and not as an internal control, all reactions were run in triplicates. As negative control, for each condition tested and each target analysed a reaction with the RNA as template was run. Reaction mixes (20 µl/reaction) were prepared as follows: 0.33 µl sample, 100 nM primers, 1x QX200 ddPCR EvaGreen supermix (Bio-Rad, 1864034). The ddPCR reactions were set up as described for the TaqMan probe-based assay above, with the only difference being that the QX200 Droplet Generation Oil for EvaGreen (Bio-Rad,1864005) was used for emulsion. For analysis, the *ZEB2* copy number values were normalized to the copies of the loading control *EIF2B2* and mean and standard error of the mean (SEM) values for the three technical replicates were calculated and reported as normalized mRNA fold change.

### RNA-seq sample preparation and sequencing

Organoids generated from 3 replicate batches of H9 and G1 cell lines were collected at 7 time points for RNA-seq analysis: 0, 2, 3, 5, 10, 15 and 25 days post neural induction. ∼300 organoids per replicate were collected at the time points ranging between day 0 and 5, ∼150 organoids at day 10, ∼100 organoids at day 15, and ∼50 organoids at day 20. Organoids were collected in Eppendorf tubes, washed with PBS, flash-frozen in liquid nitrogen and stored at −80 °C.

RNA from all 42 samples was isolated in parallel with the Direct-zol-96 RNA kit (Zymo Research, R2055), following the product’s manual. cDNA libraries were generated according to NEBNext Ultra II Directional RNA Library Prep Kit for Illumina (NEB E7760). cDNA was amplified for 12 cycles and barcoded with NEBNext Multiplex Oligos for Illumina (NEB E7600). Libraries were tested for quality using the 2100 Bioanalyzer system (Agilent) and concentrations were measured using Qubit dsDNA HS Assay Kit (Thermo Fisher, Q32851) on Qubit 3 Fluorometer (Thermo Fisher). Samples were pooled together to 15 nM concentration and sequenced with single-end 50 base mode on 3 lanes of an Illumina HiSeq 4000 instrument.

### RNA-seq data analysis

RNA-seq reads were processed using the PRAGUI pipeline (https://github.com/lmb-seq/PRAGUI). Reads were trimmed using Trim Galore! version 0.6.3_dev (https://github.com/FelixKrueger/TrimGalore) with Cutadapt version 2.4 (Martin, 2011). Default parameters were used. Read quality was analysed by FASTQC version v0.11.5 (Andrews, 2010). Human and gorilla reads were aligned to GRCh38 and Kamilah_GGO_v0 reference genomes respectively using HISAT2 (version 2.0.0-beta). GTF gene annotation files for both of the reference genomes were downloaded and filtered for a list of 16,763 genes that are annotated with identical gene names in both species. Reads were assigned to genes in the custom GTF files and counted using HTSeq version 0.11.2 (Anders et al., 2015). Read counts were then normalized to transcripts per million (TPM) within each sample and log2 transformed (log2(TPM+1)). Unbiased clustering of the samples with principal component analysis was performed using the 3000 most variable genes. Log2 transformed TPM values were converted to z-scores, (x_i_ – μ)/σ, where x_i_ is the TPM at a given time point, μ is the mean TPM of the gene across all time points and σ is the standard deviation of the TPM across all time points. Pearson correlation was performed on z-scaled mean log2 transformed TPM and on all replicates z-scaled. Hierarchical clustering was done on z-scaled means of the 3000 most variable genes of mean log2 transformed TPMs. GO term enrichment analysis was performed using the online g:Profiler software (Reimand et al., 2007)(https://biit.cs.ut.ee/gprofiler/gost).

### Differential temporal expression analysis

To assess temporal changes in gene expression between day 3, 5 and 10, the list of 16,763 genes was first filtered based on the following criteria: 1) genes were expressed at >10 TPM in at least one time point in all replicates of both species, 2) genes displayed a fold-change of >1.5 TPM between any two time points in at least one species. This resulted in a list of 3,526 genes. Log2 normalized TPMs were then z-scaled across the three time points. Next, gene list was further filtered in order to remove genes with variable expression patterns across replicates by removing genes with a squared difference >6 in either species. The squared difference value was obtained by calculating the squared difference of z-scores per time point between all three replicates and then taking the sum of this number for all time points. This list of 2,905 genes was used for time course sequencing data analysis using the TCseq package (Wu and Gu, 2020) available online (https://rdrr.io/bioc/TCseq/f/inst/doc/TCseq.pdf). Replicates were kept separate for the analysis, meaning that the input was 17,430 unique patterns of gene expression representing 2,905 genes in 2 species and 3 replicates per species. The TCSeq analysis was run using the timeclust function with the settings algo = ‘cm’, k = 10, resulting in the unsupervised soft clustering of gene expression patterns into 10 clusters with similar z-scaled temporal patterns. Replicates of 563 genes were found to be in 3 different clusters in at least one species and were removed from downstream analysis resulting in 2,342 genes (Supplementary Data 2). 22.5% of genes, 527 genes, were found to never be in the same cluster between species showing a robustly different expression pattern (Supplementary Data 4). Genes were assigned to the cluster where 2 or all of their replicates were found per species. 59 % of genes were assigned to different clusters between species. GO term enrichment analysis was performed on the genes present in each of the 10 clusters generated by TCseq per species, resulting in a total of 159 GO:BP terms found in both species, of which 85 were found to be moving between species (Supplementary Data 3). GO term analysis on the 527 genes moving robustly between species revealed 67 were linked to enriched cell morphogenesis-related terms (“cell morphogenesis”, “cell part morphogenesis”, “cellular component morphogenesis”). These genes were intersected with a list of 1,639 confirmed transcription factors (Lambert et al., 2018), revealing 8 transcription factors related to cell morphogenesis (Figure 4F).

## Author contributions

S.B-K. designed and conducted experiments, performed bioinformatics analysis, analysed data, and wrote the manuscript. S.L.G. designed and conducted experiments, analysed data, and wrote the manuscript. M.S. optimized and cultured organoids and prepared reagents. E.S.R. wrote and optimized scripts for image analysis and quantification. P.F-P. performed bioinformatics analysis. I.K. prepared ape cells for reprogramming. S.W., U.M., and G.W. provided cell lines and intellectual input. M.A.L. designed and supervised the project, performed experiments, and wrote the manuscript.

## Data availability

RNAseq data are available on NCBI GEO, accession number GSE153076.

## Acknowledgments

The authors would like to thank members of the Lancaster lab for helpful feedback and discussions. We also thank the Light Microscopy facility of the MRC Laboratory of Molecular Biology, and M. Kellner, A. Kalinka, G. Ghattaoraya, A. Crisp and T. Stevens for help with bioinformatics. The authors thank Twycross zoo, in particular Zak Shovell and Clare Ellis, for sample collection. We also thank Johannes Schindelin for the FIJI plug-in used for cell counting. This work was supported by the Medical Research Council (MC_UP_1201/9) and the European Research Council (ERC STG 757710). S.B-K. is supported by a CRUK Cancer Centre PhD Studentship. E.S.R. is supported by CCIMI, Cantab Capital Institute for Mathematics of Information and LSM, London Mathematical Society.

## Declaration of interests

The authors have no competing interests to declare.

## Supplementary Figures

**Figure S1.**
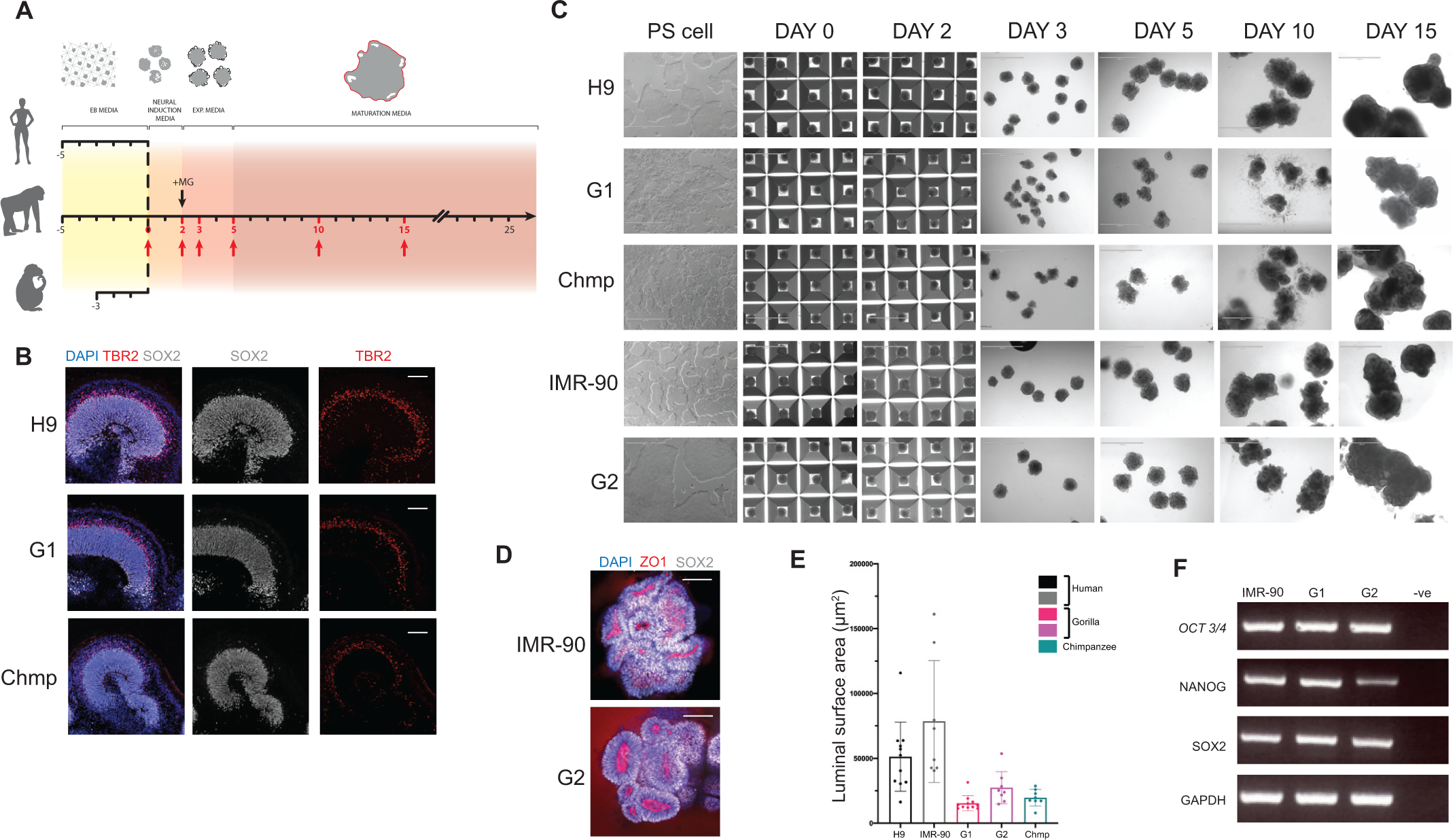
Human and ape stem cells and organoids are highly comparable in terms of identity and morphology. **A.** Schematic of the timeline for generating brain organoids in aggrewell from human, gorilla and chimpanzee stem cells. Colors represent changes in media and the various stages of the protocol and +MG represents Matrigel embedding. Note chimpanzee organoids have an EB stage that is 2 days shorter than human and gorilla. **B.** Immunofluorescent staining of representative 5-week ape organoids for neural progenitor marker SOX2 (grey), intermediate progenitor marker TBR2 (red) and DAPI (blue) show cortical lobules of similar thickness and relative proportions of cell types. Scale bar: 100 μm. **C.** Representative brightfield images of pluripotent stem cells (PS) and organoids taken throughout the differentiation protocol at the time points highlighted in red in panel A. Note the more rounded neuroepithelium observed in day 5 organoids generated from nonhuman ape (Chmp, G1 and G2) cells versus day 5 organoids generated from human (H9, IMR-90) cells. Scale bar: 1 mm. **D.** Representative immunofluorescence images of the center of whole mount human (IMR-90) and gorilla (G2) day 5 organoids with staining for ZO1 and SOX2 showing polarized neural progenitor cells organized around rounded (gorilla) and more convoluted (human) ZO1 positive apical lumens. Scale bar: 100 μm. **E.** Quantification of the surface area of the largest apical lumen per day 5 organoid showing calculations for organoids derived from additional human (IMR-90) and gorilla (G2) cell lines. The data reveals the same trend of more expanded buds in human versus non-human ape. Mean luminal surface area: IMR-90 = 78,463 μm^2^; G2 = 27,426 μm^2^; H9, G1, Chmp are reported in Figure 1F. **F.** RT-PCR shows expression of pluripotency markers (*OCT3/4, NANOG, SOX2*) and *GAPDH* loading control in IMR-90, G1 and G2 cell lines. –ve is the water negative control.

**Figure S2.**
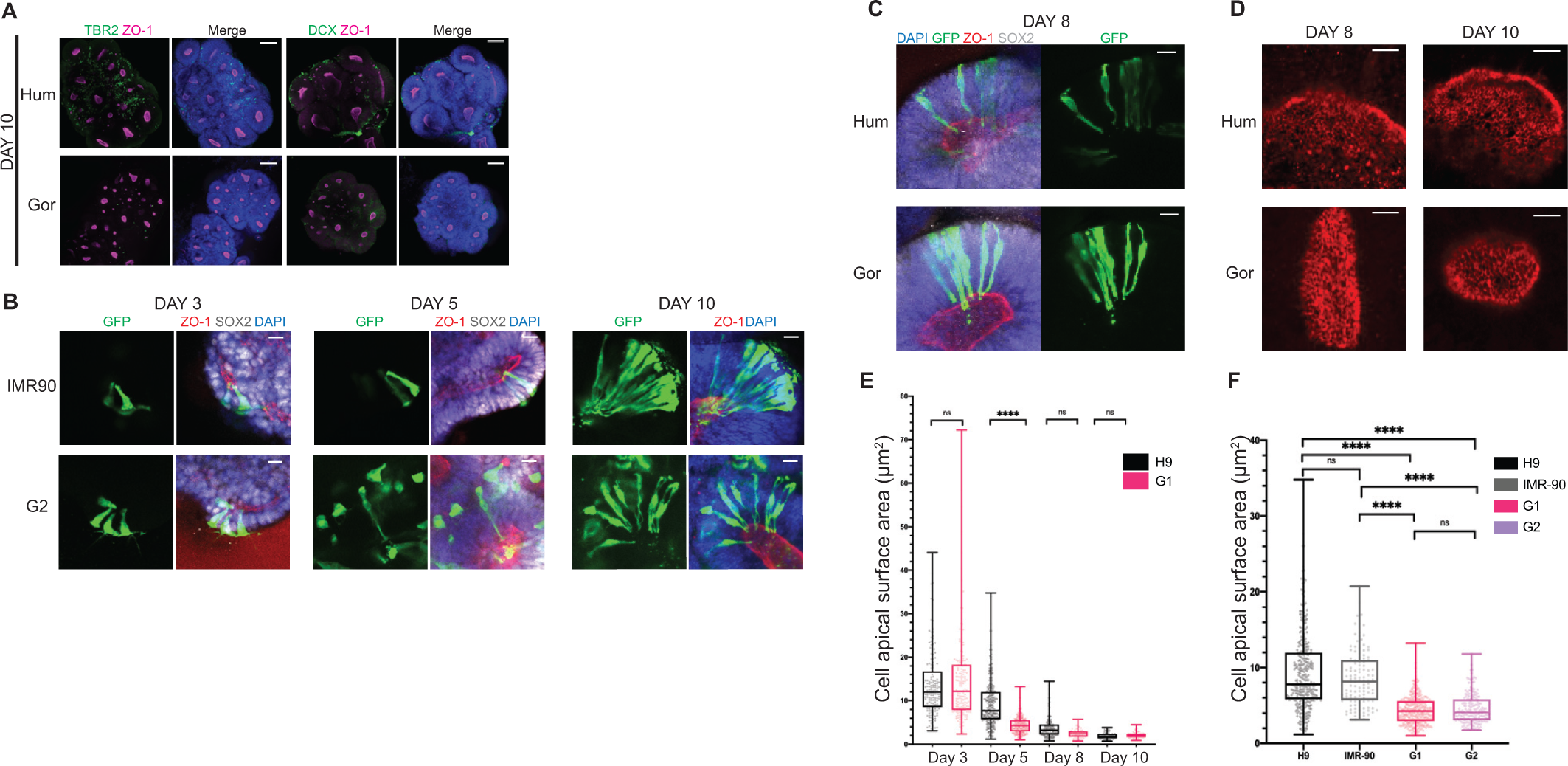
Ape NE cells undergo cell shape transition before the onset of neurogenesis. **A.** Immunofluorescent staining for newborn neurons (DCX) and intermediate progenitors (TBR2) at day 10 human (H9) and gorilla (G1) organoids. Note the sparse presence of these cell types, indicating neurogenesis is only just beginning. DAPI is shown in blue. Scale bar: 100 μm. **B.** Representative immunofluorescence images stained for GFP, ZO1 and SOX2 showing the morphology (GFP+) of neural progenitor cells in organoids derived from additional human (IMR-90) and gorilla (G2) cell lines. Progenitor cell shapes at day 3, 5 and 10 observed in organoids derived from IMR-90 and G2 reflect the shape changes seen in H9 and G1 derived organoids (Figure 2C-D) respectively. Both cell lines exhibit columnar NE and RG morphologies at day 3 and day 10 respectively, while at day 5 IMR-90 cells exhibit columnar and G2 cells exhibit tNE morphologies. Scale bar: 20 μm. **C.** Representative immunofluorescence images of sparsely labeled progenitor cells (SOX2+) at day 8 show thinning of apicobasal processes and tNE morphologies in both species (human H9, gorilla G1). Scale bar: 20 μm. **D.** Immunofluorescent staining for ZO1 on the surface of apical lumens at day 8 and 15 showing constricted apical surfaces of individual progenitor cells in both species. Scale bar: 10 μm. **E.** Quantification of the surface area of individual human (H9) and gorilla (G1) neural progenitor cells between day 3 and 10 shows a gradual reduction in apical surface area over time in both species by 7-fold. Mean apical surface area/cell: human day 3 = 13.82μm^2^; gorilla day 3 = 14.80μm^2^; human day 5 = 9.39μm^2^; gorilla day 5 = 4.48μm^2^; human day 8 = 3.86 μm^2^; gorilla day 8 = 2.46 μm^2^; human day 10 = 1.92μm^2^; gorilla day 10 = 2.13μm^2^. *P<0.05, **** P<0.0001, Kruskal-Wallis and post-hoc Dunn’s multiple comparisons test, n (day 3, human) = 164 cells from 8 organoids and 2 batches, n (day 3, gorilla) = 176 cells from 8 organoids and 2 batches, n (day 5) = reported in Figure 2G, n (day 8, human) = 171 cells from 4 organoids, n (day 8, gorilla) = 55 cells from 2 organoids, n (day 10, human) = 68 cells from 3 organoids, n (day 10, gorilla) = 74 cells from 4 organoids. Error bars are min-max values, dots on the boxplot represent individual cells. **F.** Day 5 quantification of apical surface area of progenitor cells showing gorilla cells are more apically constricted than human in organoids derived from additional human (IMR-90) and gorilla (G2) cell lines. Mean apical surface area/cell: IMR-90 = 8.53 μm^2^; G2 = 4.54 μm^2^; H9, G1 reported in Figure 2G. *P<0.05, **** P<0.0001, Kruskal-Wallis and post-hoc Dunn’s multiple comparisons test, n (H9, G1) = reported in Figure 2G, n (IMR-90) = 121 cells from 4 organoids, n (G2) = 172 cells from 5 organoids. Error bars are min-max values, dots on the boxplot represent individual cells.

**Figure S3.**
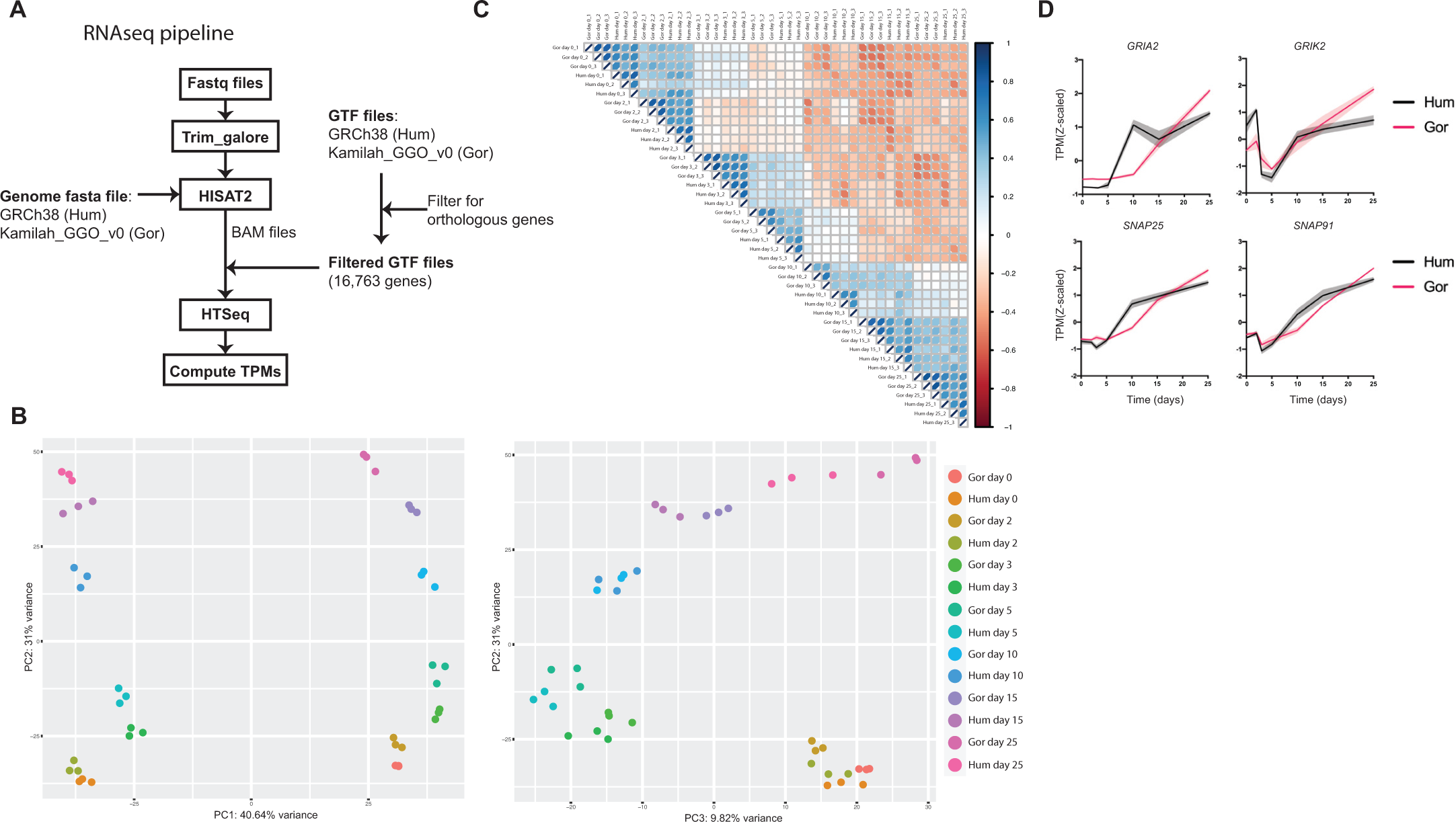
RNAseq data analysis pipeline and normalization. **A.** Workflow summarizing the RNAseq analysis pipeline (see methods). **B.** PCA performed using log2-transformed TPMs. Samples are color-coded by time point and species. Graph of PC1 vs PC2 (left) shows biological replicates grouping together, samples separating by species along PC1 and separating by time point along PC2. Plotting PC2 vs PC3 (right) shows samples separating by time point and not species. **C.** Pearson’s correlation of all samples using z-scaled log2-transformed TPMs of all genes. Darker blue depicts stronger positive correlation between samples and darker orange a stronger negative correlation. Data shows the strongest correlation between biological replicates within a species followed by between species time point-matched samples. Correlation between species is lowest at day 5 and 10. **D.** Temporal expression pattern (z-scaled) of genes related to synaptic formation and maturation (*GRIA2, GRIK2, SNAP25, SNAP91*). Shaded error bar is S.D.

**Figure S4.**
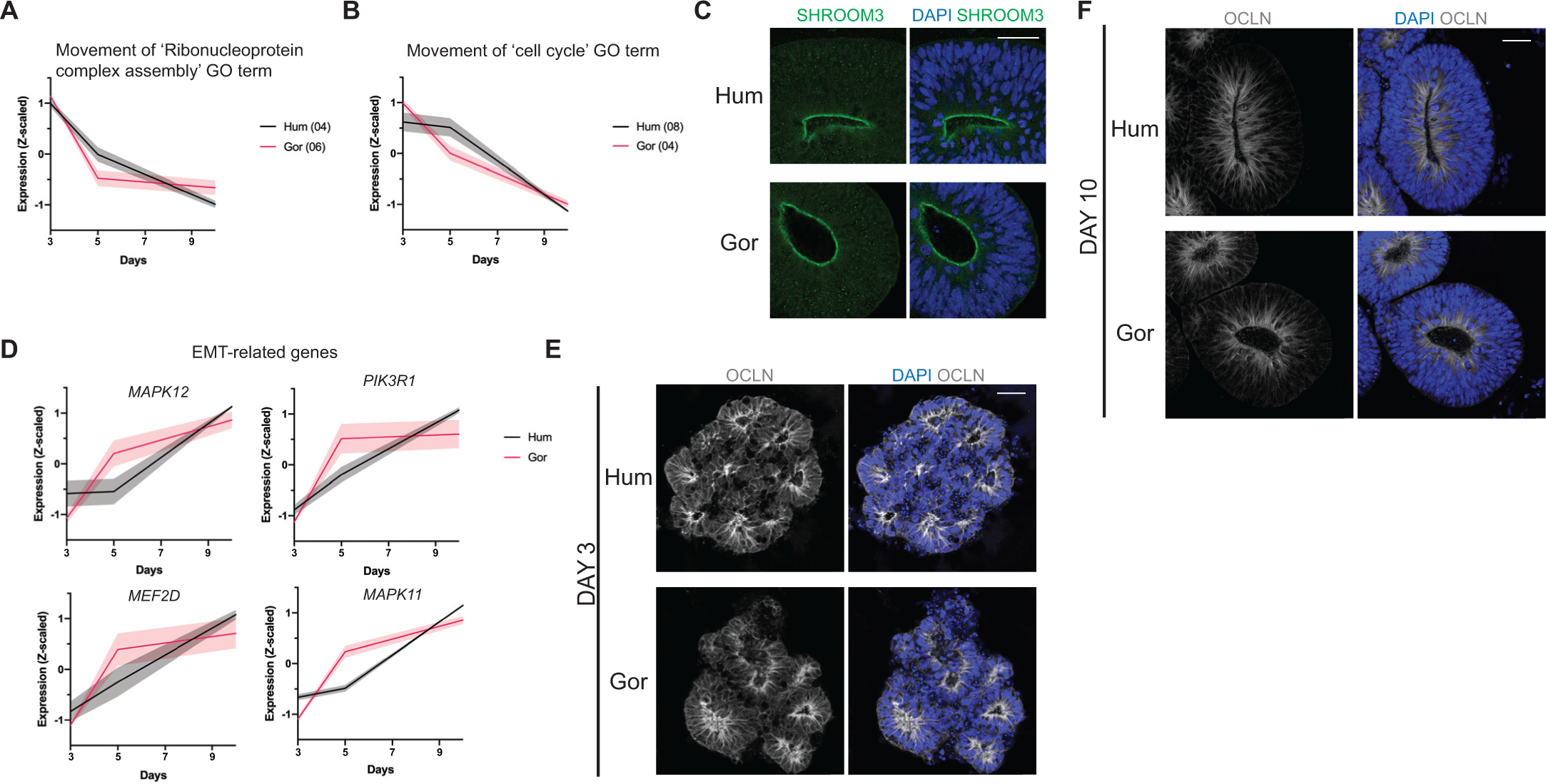
Expression patterns of key factors with differential temporal dynamics. **A, B.** Mean temporal expression pattern (z-scaled) of genes in clusters enriched for: **A.** ‘Ribonucleoprotein complex assembly’-related GO:BP terms (human cluster 4, gorilla cluster 6) **B.** ‘cell cycle’-related GO:BP terms (human cluster 8, gorilla cluster 4)**. C.** Immunofluorescent staining of day 10 organoids for SHROOM3 shows strong apical expression in neuroepithelium of both species (H9, G1). Scale bar: 40 μm. **D.** Mean temporal expression (z-scaled) of EMT-related genes enriched for WP term ‘Epithelial to mesenchymal transition in colorectal cancer’ *MAPK12, MEF2D, PIK3R1* and *MAPK11*, robustly changing pattern between species in the same way enrichment of ‘cell morphogenesis’-related terms changes pattern between species (Figure 4B). **E,F.** Immunofluorescent staining of organoids (H9, G1) for OCLN at **E.** day 3, showing expression along the apicobasal length of progenitor cells in both species **F.** day 10, showing lowered expression limited apically in both species. Scale bar: 40 μm.

**Figure S5.**
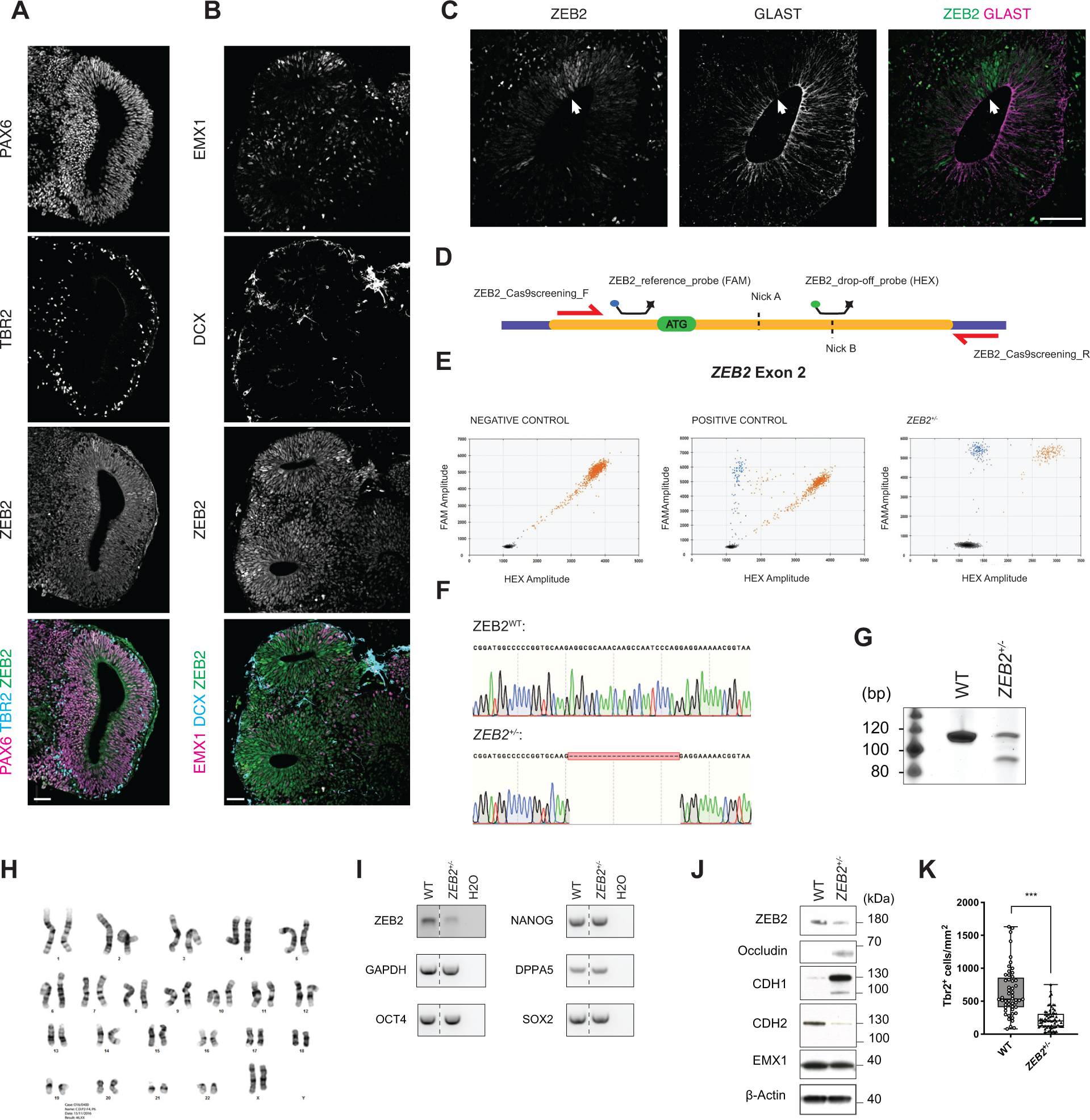
ZEB2 expression and targeting for loss of function. **A.** Representative immunofluorescence image of a day 16 human (H9) organoid showing ZEB2 expression in PAX6+ progenitor cells of cortical lobules with dorsal telencephalic identity, as indicated by TBR2+ intermediate progenitor cells. Scale bar: 50 µm. **B.** Representative immunofluorescence image of a day 16 human (H9) organoid showing ZEB2 expression in EMX1+ dorsal telencephalic progenitors of early neurogenic organoids, as marked by the sparse DCX^+^ staining. Scale bar: 50 µm. **C.** Representative immunofluorescence image of a day 25 human (H9) organoid showing a mutually exclusive pattern of expression between ZEB2 and the radial glia marker protein GLAST. Scale bar: 100 µm. **D.** Schematic representation of the CRISPR-Cas9n editing strategy, where the first coding exon of the *ZEB2* gene (exon 2, NCBI ref sequence NM_014795.4:182-323) was targeted by two nickases (dashed lines) and screening was performed by assaying the drop-off frequency of a HEX-labelled probe, binding to one of the nick sites, relative to a FAM-labelled reference probe binding away from the disrupted region. The exon is marked in orange while introns are marked in purple. **E.** Example ddPCR 2D scatter-plots of a negative control sample (HEK293 cells), showing only a FAM-HEX double positive (red) and an empty droplet cluster (black) and a positive control sample (HEK293 cells expressing WT Cas9 and *ZEB2* guides), showing a FAM-only cluster (blue) in the upper-left quadrant of the 2D plot corresponding to edited alleles. ddPCR 2D scatter-plot of the H9 ZEB2^+/−^ hESC edited line showing a 1:1 ratio between the WT and edited allele. **F**. Representative chromatograms of the *ZEB2* alleles in the H9 *ZEB2*^+/−^ hESC cells. The CRISPR-Cas9 target region was PCR amplified with a high fidelity polymerase, the PCR product was blunt-end cloned into the pJET1.2 vector and following purification, plasmids from different colonies carrying the insert were sequenced. Sequencing reveals that the edited allele harbors a 23 bp deletion. **G**. DNA-PAGE analysis of a short PCR amplicon spanning the CRISPR-Cas9 *ZEB2* target site in WT H9 and H9 ZEB2^+/−^ hESCs. The gel reveals the presence of two bands, corresponding to the WT and the edited allele in H9 ZEB2^+/−^ hESCs. **H.** Representative images of karyotype analysis on 20 G-banded metaphase spreads from the H9 ZEB2^+/−^ hESCs used to generate the stock. The cell line displays normal karyotype. **I.** RT-PCR analysis for expression of *ZEB2*, the key pluripotency markers *SOX2, NANOG, OCT4* and *DPPA5* and the loading control *GAPDH*. PCR shows that upon a ∼50% reduction in *ZEB2* mRNA levels the mutant stem cells retain expression of pluripotency markers at comparable levels to WT H9 hESCs. WT and *ZEB2*^+/−^ were run on the same gel but not adjacent to each other, the dashed line indicates where the gel was spliced. **J.** Western blot of H9 WT and ZEB2^+/−^ organoids at day 16 for ZEB2, the tight-junction protein Occludin, the adherens junction components CDH1 and CDH2, the dorsal telencephalic marker EMX1 and the loading control β-Actin. The blots show a sizeable increase in CDH1 and Occludin and a decrease in CDH2 whilst EMX1 levels, and thus dorsal telencephalic identity appears to be largely unaffected. (K) Box and whiskers plot reporting the quantifications of the number of TBR2+ cells (TBR2+ cells/mm^2^) in day 16 WT and *ZEB2*^+/−^ organoids. Quantifications were performed on n=52 WT and n=68 *ZEB2*^+/−^ ventricles corresponding to 12 organoids from 2 distinct batches. A Mann-Whitney U test was used for statistical comparison (p<0.0001).

**Figure S6.**
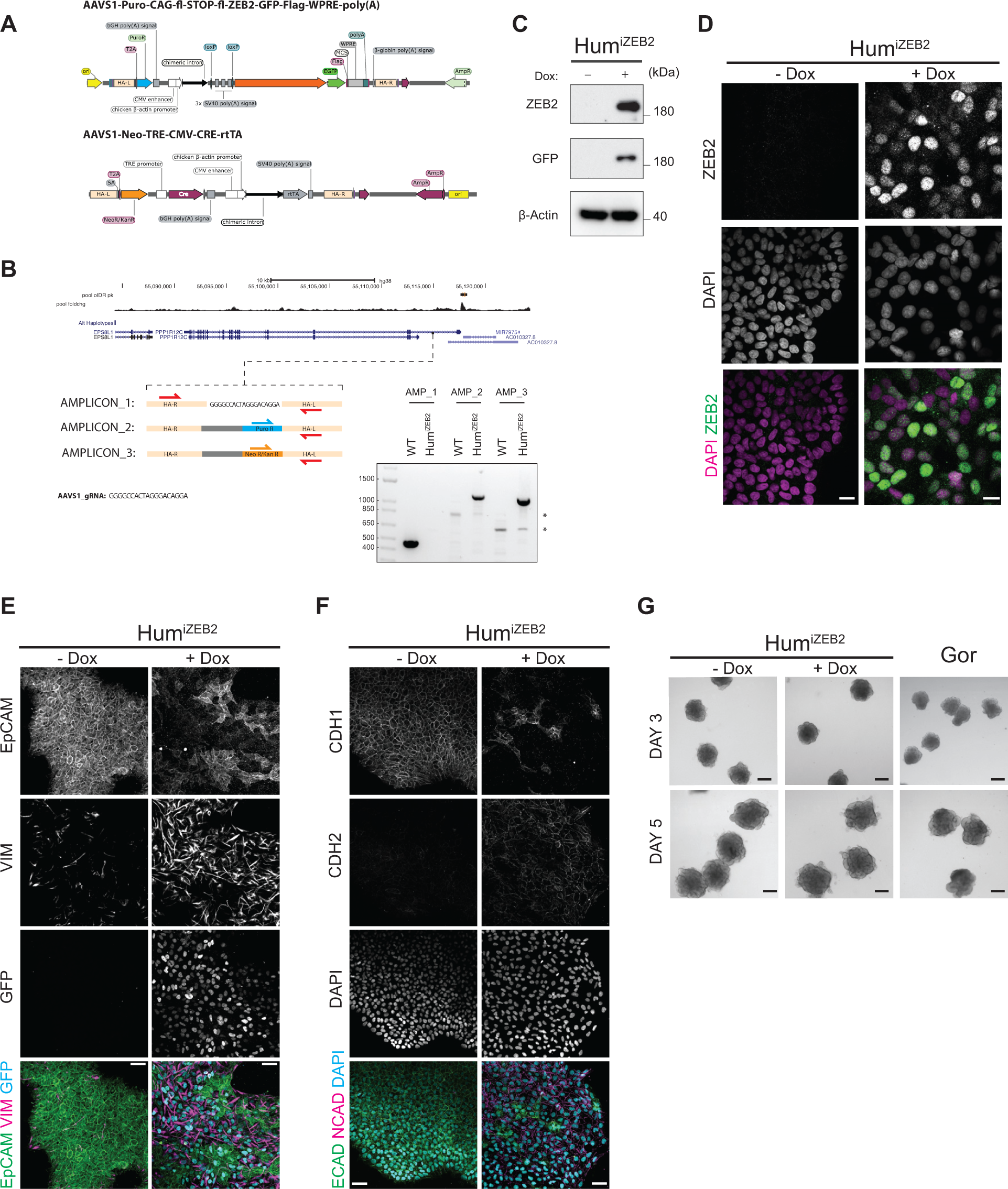
Early forced expression of ZEB2 in human NE cells. **A.** Plasmid maps of the CRISPR homology-directed repair (HDR) templates used to target the AAVS1 safe-harbour locus in H9 hESC cells – top is the CAG-lox-STOP-lox-ZEB2-GFP-Flag inducible expression construct and bottom is the construct encoding CRE recombinase under the control of a tetracycline responsive promoter and the reverse tetracycline transactivator (rtTA) driven by the CAG promoter. **B**. UCSC Genome Browser view of the AAVS1 locus and CRISPR-Cas9 targeting strategy of intron 1 of *PPP1R12C*. The promoter-less splice-acceptor (SA), T2A peptide-linked “gene trap” is such that expression of the promoter-less selection cassette is driven by the endogenous *PPP1R12C* gene, thus effectively eliminating false-positive background arising from random integration. The panel reports the PCR genotyping strategy – upon successful targeting of the AAVS1 locus, while amplicon 1 is lost due to the size increase following insert integration, amplicons 2 and 3 are gained. The PCR gel shows successful genotyping of the colony used for all experiments shown. The asterisks mark unspecific bands **C**. Transgene induction test in H9^iZEB2^ cells treated with and without doxycycline for 6 days and assayed by western blot for ZEB2, GFP and β-actin. **D**. Immunofluorescence images of 6-day induced and uninduced H9^iZEB2^ cells stained for ZEB2 and DAPI, showing that doxycyline induction results in ZEB2 expression and nuclear translocation, without adverse effects on its localisation due to tagging with GFP. Scale bar: 20 µm **E**. Immunofluorescence images of 6-day induced and uninduced H9^iZEB2^ cells stained for GFP, Vimentin and EpCAM. The data reveal a reduction in EpCAM expression and an increase in Vimentin expression following expression of ZEB2-GFP. Scale bar: 50 µm **F**. Immunofluorescence images of 6-day induced and uninduced H9^iZEB2^ cells stained for DAPI, CDH1 and CDH2. The data reveal a reduction in CDH1 expression and an increase in CDH2 expression following induction. Scale bar: 50 µm. **G**. Brightfield images of induced (+ Dox) and uninduced (-Dox) H9^iZEB2^ organoids and gorilla (G1) organoids at day 3 and day 5, showing indistinguishable tissue architecture between organoids at day 3, while day 5 organoids show neural buds that are more rounded in gorilla and ZEB2 induced vs uninduced organoids. Scale bar: 200 μm.

**Supplementary Table 1.**
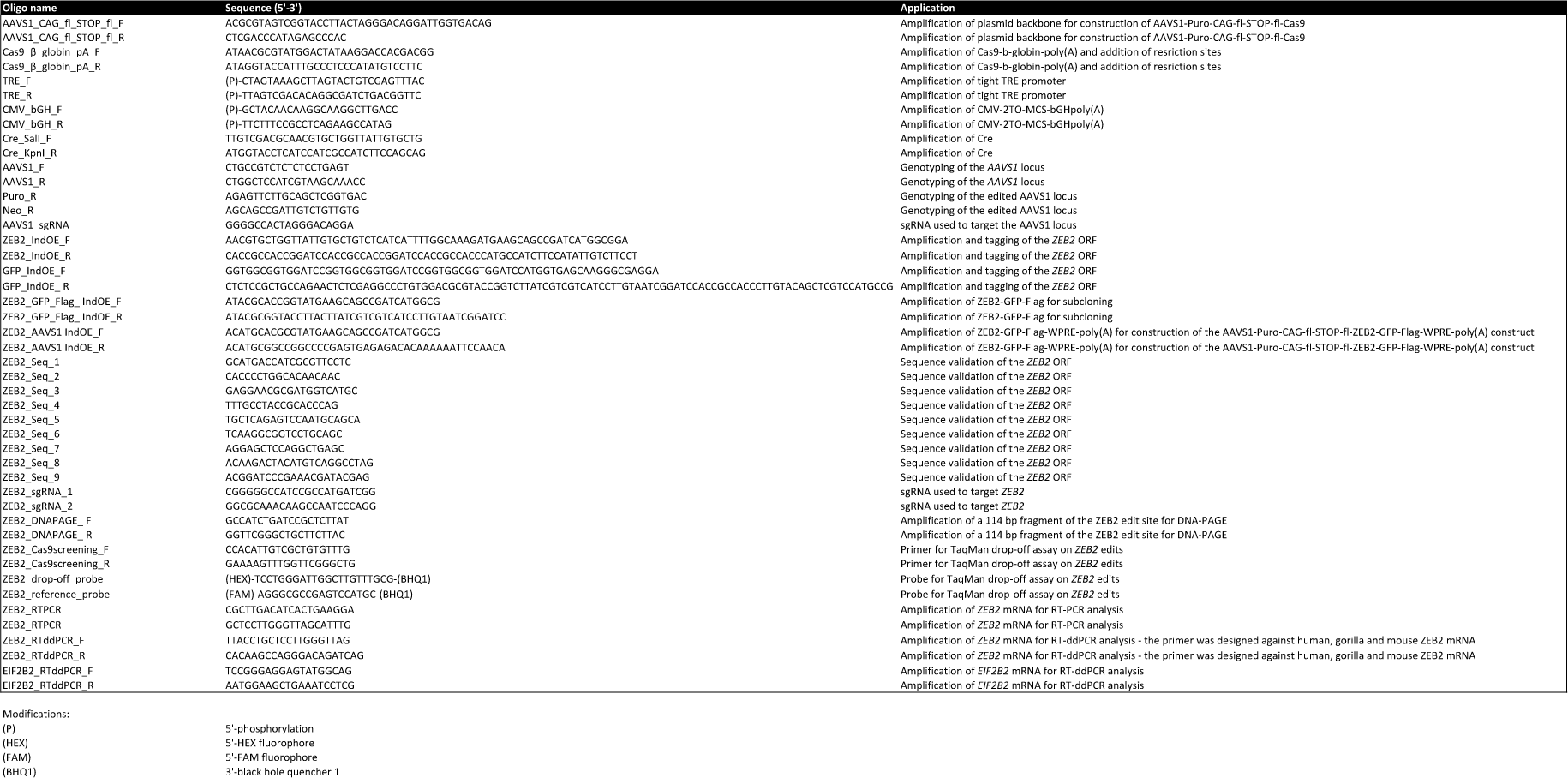
A list of primers and DNA oligos employed in the study for cloning and genome editing.

**Movie S1.** Live imaging of progenitor cells sparsely labelled with GFP in a human cerebral organoid beginning on day 3. Images were acquired every 20 minutes. Note the appearance of columnar neuroepithelial cells at the onset of the video with apical division (06:20 bottom-right is apical) with a loss of basal processes during division. 7 hours post division (the last frame) shows apical processes that have gradually re-thickened in human since division. Scale bar: 100 μm.

**Movie S2.** Live imaging of progenitor cells sparsely labelled with GFP in a gorilla cerebral organoid beginning on day 3. Images were acquired every 20 minutes. Note the appearance of columnar neuroepithelial cells at the onset of the video with apical division (05:00 right is apical) with a loss of basal processes during division. 7 hours post division (the last frame) shows apical processes that have remained thin in the gorilla. Scale bar: 100 μm.

**Supplementary Data 1.** Related to figure 3B and 3F. List of top 300 variable genes at 5, 10, and 25 days and the corresponding GO term enrichment tables generated from g:Profiler (Reimand et al., 2007).

**Supplementary Data 2.** List of the 2,342 genes used for time course analysis using TCseq, and their assigned clusters. Each gene is also annotated by species, human (Hum) or gorilla (Gor).

**Supplementary Data 3.** Related to figure 4A. A list of the GO:BP terms that were found to be enriched in TCseq clusters and the corresponding species and cluster.

**Supplementary Data 4.** List of the 527 genes that shifted robustly between different clusters of each species and the corresponding clusters.

## Notes

### Competing Interest Statement

The authors have declared no competing interest.

